# Proteolytic control of the SARS-CoV-2 furin cleavage site defines the phenotypic evolution from the pandemic to endemic state

**DOI:** 10.64898/2026.09.23.753140

**Authors:** Anupriya Aggarwal, Samantha Ognenovska, Timothy Ison, Christina Fichter, Vanessa Milogiannakis, Madeeha Afzal, Alberto Ospina Stella, Daniel Asarnow, Angelica L. Morgan, Shafagh Waters, Till Boecking, Boaz Ng, Camille Esneau, Nathan Bartlett, Atharva Naik, Stefan Pöhlmann, Markus Hoffmann, Stefan Bröer, Angelika Bröer, Matt D Johansen, Robert Brink, Louise M Burrell, Sheila K Patel, Sally Ellis, Michael Wehrhahn, Elena Martinez, Andrew Ginn, Melissa J Churchill, Thomas A Angelovich, William Rawlinson, Malinna Yeang, Greg Walker, Charles Foster, Jen Kok, Vitali Sintchenkov, Rebecca Rocket, William Asquith, Rhys Parry, Julian D Sng, Greg Neely, Cesar Moreno, Lipin Loo, Mathew Spence, Colin Jackson, Anthony D Kelleher, Young-Jun Park, Cecily Gibson, Ben Merz, Cameron Stewart, Fabienne Brilot, Alexander A Khromykh, David Veesler, Vineet D Menachery, Stuart G Turville

**Author notes:** Equal author contribution (Anupriya Aggarwal, Samantha Ognenovska, Timothy Ison).

## Abstract

The successive emergence of SARS-CoV-2 variants with altered tissue tropism has progressively decoupled transmissibility from lower respiratory tract pathogenicity. Omicron lineages transmit exceptionally well whilst limiting severe lung disease. Here, we demonstrate that this phenotype has evolved through two temporally distinct tissue-specific proteolytic controls at the Spike (S) furin cleavage site (FCS). At the virion level, cell-type-dependent furin-mediated FCS hyper-cleavage depletes S in lung but not nasal epithelial cells. At the infected cell membrane during cell-cell spread, TMPRSS2 mediates S FCS cleavage but is negatively regulated in the presence of ACE2. Tissue-specific solute carriers SLC6A19 and SLC6A20 sequester ACE2, thereby relieving inhibition of TMPRSS2 and enabling S FCS cleavage during cell-cell spread. Critically, whilst all SARS-CoV-2 lineages benefit from this latter pathway, Omicron variants have evolved exclusive dependence on it—a strategic consolidation that focuses S proteolytic activation to TMPRSS2 alone. The combined outcomes of S protein regulation across viral and cell membranes reveal how tissue-specific proteolytic optimisation drives Omicron’s transmission fitness advantage in the upper respiratory tract but at the cost of heavy attenuation in the lower respiratory tract.

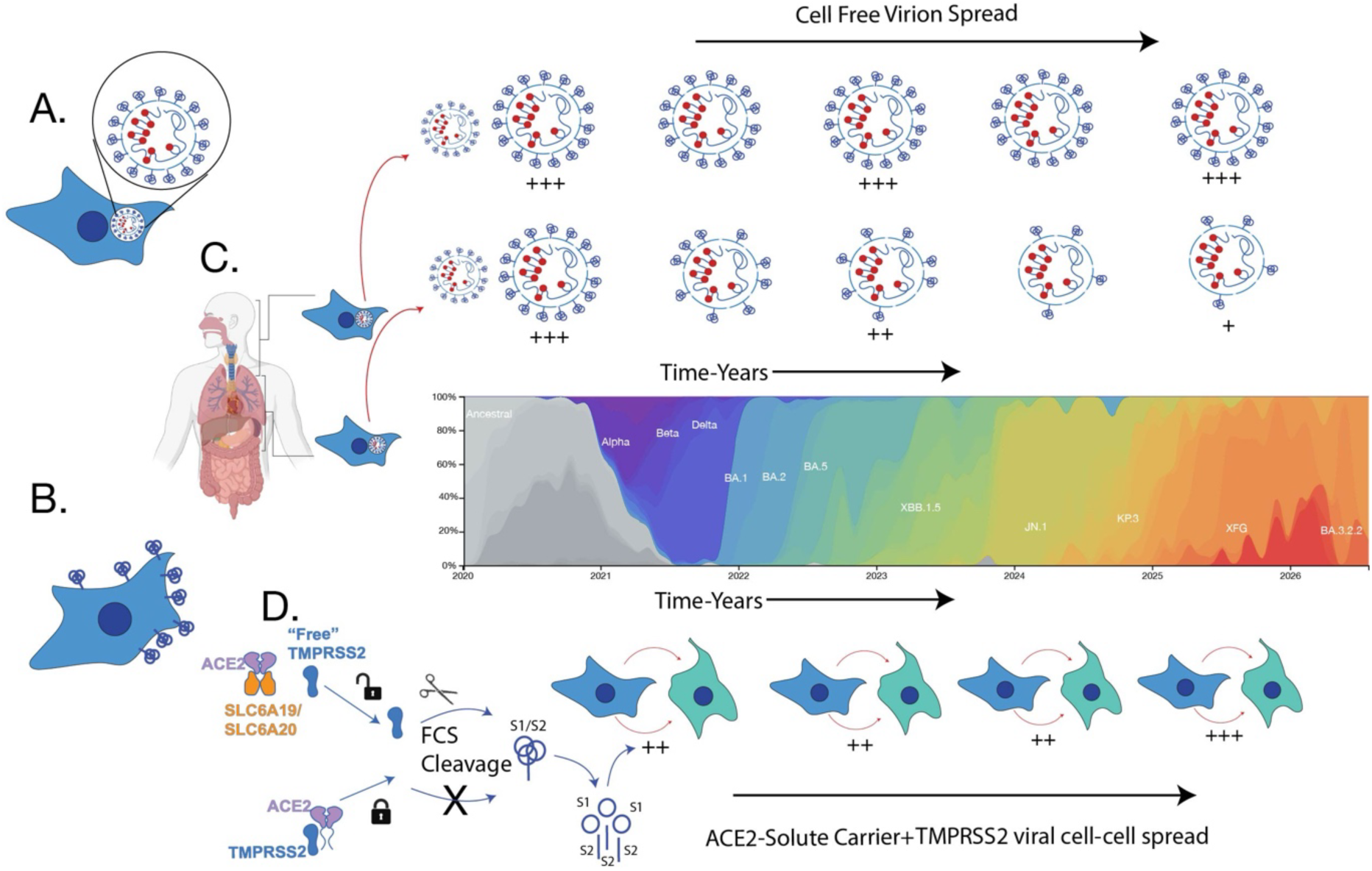

**Graphical Abstract: Dynamics of Spike pools, S FCS cleavage and ACE2 conformations collectively define Omicron Tropism**

**A & B. Spatial regulation of S protein FCS cleavage.** Viral S protein can traffic to two distinct cellular locations: **A.** virion particles within the trans-Golgi network of infected cells, or **B.** the plasma membrane of infected cells. Proteolytic cleavage of the FCS at each location is governed by different mechanisms. **C.** In virion particles, furin plays the dominant role, mediating hyper-FCS cleavage that culminates in S protein depletion in virions generated from cells derived from the lung. Nasal epithelia and other cells permissive to Omicron infections, sustain efficient FCS cleavage without S protein virion depletion. Over time, this phenotype has continued to consolidate from the pandemic to endemic stages of SARS-CoV-2 (Spike depletion relative to Global frequencies of major variants (courtesy of Nextstrain)). The arrival of JN.1 sub-lineages observed the peak of S protein depletion from furin and furin like protease hyper-cleavage. **D.** From the consequences of S protein depletion, the role of non-FCS cleaved S at the infected cell membrane is revealed. At the infected cell membrane (cells in blue) during cell-cell contact, TMPRSS2 is the primary FCS protease and is present on the recipient cell membrane (cells in green). mportantly, ACE2 negatively regulates TMPRSS2 activity at this location; however, this inhibition is circumvented when ACE2 forms complexes with solute carriers SLC6A19 or SLC6A20. This interaction frees TMPRSS2 from ACE2-mediated inhibition, enabling efficient S1/S2 FCS cleavage at the cell membrane by TMPRSS2. This subsequently exposes S2’ for TMPRSS2 cleavage, driving efficient cell-cell spread via a common protease-gated mechanism. Efficient cell-cell spread is sustained across all SARS-CoV-2 lineages using this pathway, and more recently demonstrated to afford JN.1 sub-lineages a fitness advantage relative to XBB.1.5 sub-lineages.

## Introduction

The four endemic coronaviruses (229E, OC43, NL63, and HKU1) which circulate globally cause mild to moderate upper respiratory tract infections. In contrast, severe acute respiratory syndrome coronavirus (SARS-CoV-1), Middle East respiratory syndrome coronavirus (MERS-CoV) and pre-Omicron variants of SARS-CoV-2 are able to infect the lower respiratory tract and cause severe respiratory disease. However, SARS-CoV-2 Omicron sub-lineages appear to have partly lost this ability and more frequently cause mild disease. A better understanding of the coronavirus cell and organ tropism is required to resolve SARS-CoV-2 pathogenesis and its phenotypic trajectory over the course of the pandemic to endemic stages.

Early SARS-CoV-2 lineages used the cellular protein angiotensin-converting enzyme 2 (ACE2) as a primary receptor, alongside the cellular subtilisin-like proprotein convertase protease furin and the transmembrane serine protease 2 (TMPRSS2) for S protein activation. SARS-CoV-2 S activation starts with cleavage at the S_1_/S_2_ furin cleavage site (FCS) during S protein transit through the secretory pathway of infected cells. Subsequent TMPRSS2-mediated cleavage at the S_2_’ site completes S activation and culminates in membrane fusion. Notably, TMPRSS2 cleavage at the S_2_’ site is promoted by prior cleavage of the FCS at S_1_/S_2_ ^4–6^. Although TMPRSS2 is classically regarded as an S_2_’ protease, biochemical studies have demonstrated that TMPRSS2 can substitute for furin and also cleave the canonical S_1_/S_2_ FCS site in the context of ectodomain trimers^3^. Furthermore, TMPRSS2-dependent, furin-independent FCS cleavage has also been observed during SARS-CoV-2 cell-cell spread ^7^ but its contribution to viral fitness across variants is largely unknown.

The evolution of the first variants of concern (VOC), Alpha, Beta, Gamma and Delta, were driven by S mutations that provided a fitness gain and promoted humoral immune evasion. Fitness mutations such as N501Y ^8,9^ increased affinity to ACE2 in Alpha, Beta, & Gamma, but were outcompeted by Delta, through augmented furin-mediated cleavage and cell entry by acquisition of the P681R within the FCS^10–15^. The subsequent rapid replacement of the Delta variant by the Omicron BA.1/BA.2 variants in late 2021 constituted a major shift in SARS-CoV-2 evolution. These variants harboured an unusually high number of mutations in the S protein that allowed for an unprecedented level of antibody evasion^16–18^. In parallel, the mutations shifted the viral tropism from the lower to the upper respiratory tract^2,13,14,19–21^, allowing for high levels of transmission through lower mean generation intervals ^22^. This fundamental shift in tropism remains unexplained but has contributed to the course of the pandemic with respect to clinical presentation of COVID-19. The leading hypothesis proposed that Omicron’s tissue tropism shift resulted from limited S FCS processing, causing a shift from TMPRSS2-mediated plasma membrane fusion to cathepsin-dependent endosomal fusion. This model was supported across multiple independent studies^23^ and appeared to explain Omicron’s altered entry pathway. However, this hypothesis does not fully account for Omicron’s continued transmission fitness and progressive genotypic changes in and around the FCS, suggesting an alternative mechanism rooted in FCS optimisation rather than cleavage resistance.

Using clinical viral isolates spanning 2020–2026, we identified cell-type-dependent hyper-cleavage of the S protein FCS as the first step towards Omicron tropism, acting by depleting S from viral particles and blocking cell-free spread. This S depletion then reveals a second phenotypic shift in Omicron lineages that proceeds during cell-cell spread: Exclusive use of TMPRSS2 for S FCS cleavage at the infected cell membrane. The latter is modulated by tissue-specific ACE2 ultra-structural organization defined by complex formation with one of two solute carrier proteins: SLC6A19 in the gut and SLC6A20 in the respiratory tract. Although S protein trafficking and regulation vary across viral and infected cell membranes, S FCS cleavage regulation is central to both pathways and culminates in a final phenotype that has defined the pandemic to endemic shift primarily caused by consolidated tropism to the upper respiratory tract.

## Results

### The Omicron Paradox: Attenuation from co-expression of the known entry factors ACE2 & TMPRSS2

To resolve the entry requirements across both pre-Omicron and Omicron lineages, we initially engineered cell lines with known levels of SARS-CoV-2 entry factors ACE2 and TMPRSS2. ACE2 and TMPRSS2 co-expressed at similarly high levels (VeroE6-ACE2-TMPRSS2 cell line) observed augmented infection by all pre-Omicron variants, but in contrast generated a continuum of attenuation in Omicron sub-lineages (Fig. 1 A-D). In contrast, expression of high TMPRSS2 alongside low endogenous levels of ACE2 sustained similar replication outcomes across pre-Omicron and Omicron lineages (Fig. 1 A-D).

**Figure 1.**
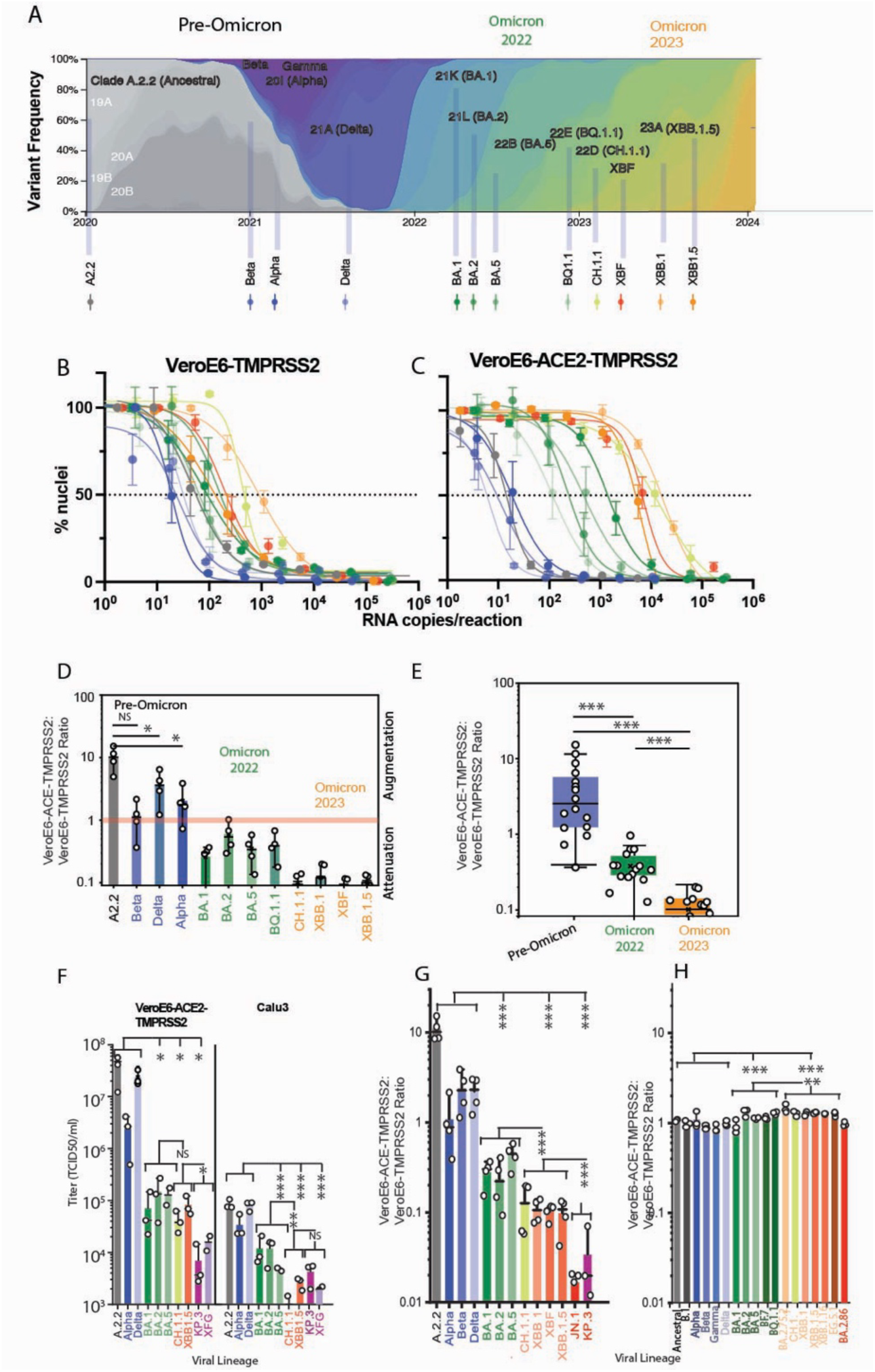
ACE2 and TMPRSS2 co-expression attenuates Omicron lineage use of TMPRSS2. **A.** Frequency of SARS-CoV-2 variants over time (nextstrain.org/sars-cov-2), with representative SARS-CoV-2 isolates selected for testing. **B&C.** Viral replication is enumerated through the accumulation of cytopathic effects that in turn lead to dose dependent cell loss (% Nuclei counts). **B.** VeroE6-TMPRSS2 cell line and **C.** VeroE6-ACE2-TMPRSS2 cell line. Left to right shifts in the dose-dependent sigmoidal curves highlight attenuation of viral replication in C. as Omicron lineages appear. Error bars represent standard deviations from 8 technical replicates. Titration curves representative of greater than 5 independent experiments. **D.** Replication of variants in the VeroE6-ACE2-TMPRSS2 line was compared relative to the parental VeroE6-TMPRSS2. (VeroE6-ACE2-TMPRSS2)/ VeroE6-TMPRSS2 titer establishes levels of augmentation or attenuation across variant eras. The red line denotes where titers of VeroE6-ACE2-TMPRSS2 are equivalent relative to its VeroE6-TMPRSS2 parent. Error bars are standard deviations across independent experiments. Each point represents an independent experiment. NS = Not significant, *P<0.05 (Dunnett’s multiple comparisons test; Supplementary Table SI). **E.** (VeroE6-ACE2-TMPRSS2)/ VeroE6-TMPRSS2 titer groupings based on Omicron Lineage period (early 2022 Omicron versus later BA.2.75 derived 2023 sub-lineages. Points derived from independent experiments in E. Error bars represent standard deviations. ***P<0.0001 (Mann Whitney U test; Supplementary Table SI). **F.** Validation of results in the engineered VeroE6-ACE2-TMPRSS2 line versus the unmodified lung-derived Calu3 cell lines with equivalent viral RNA copies in each expanded isolate (all experiments n=3), **p<0.05, **p<0.001, ***p<0.0001; Mann Whitney U test, Supplementary Table SI). **G.&H.** Cross validation of primary isolate infections with Pseudotyping of S can determine if attenuation is at the level of entry. For both G. & H., ratios are established by use of VeroE6-ACE2-TMPRSS2 relative to its VeroE6-TMPRSS2 parent. Error bars are standard deviations across independent experiments. Each point represents an independent experiment. **G.** Infection outcomes using primary isolates **H.** Contrasting outcomes using S Pseudotyping. *P<0.01, **P<0.001, ***P<0.0001 (Mann Whitney U test; Supplementary Table SI).

For the VeroE6-ACE2-TMPRSS2 line, peak augmentation was observed in the earliest circulating clade A lineage with the S protein identical to early Clade A lineage initially detected in Wuhan at the start of the pandemic (Fig. 1 E). In contrast, Omicron lineage attenuation appeared initially across two generations. The first generation were marked by early Omicron sub-lineages BA.1, BA,2 and BA.5 (Fig. 1E & F). The second generation was based on the divergent BA.2.75 derived sub-lineages (Fig. 1E & F). Cross-validation of outcomes in the VeroE6-ACE2-TMPRSS2 with the lung-derived Calu3 cell line showed similar decreases in infectivity of SARS-CoV-2 variants as they appear over time (Fig. 1G). To determine if Omicron attenuation occurred at entry, we performed S pseudotyping assays using the same clonal VeroE6-ACE2-TMPRSS2 line. In contrast to viral replication outcomes of primary isolates (Fig. 1H), pseudotyped viruses entered cells with similar efficiency in the presence of ACE2 and TMPRSS2 (Fig. 1I), indicating that Omicron attenuation under these conditions is not at the level of entry. We concluded that ACE2-TMPRSS2 co-expression at equivalent levels negatively influenced Omicron replication at a stage occurring after entry into cells, which could thus only be observed during multi-round viral infections.

### Cell-type-dependent furin-like protease hyper-cleavage depletes S protein from Omicron virions

Omicron attenuation in ACE2-TMPRSS2 cells could not be captured by pseudotyping, implying a defect in multi-round replication rather than entry. We thus examined S protein on viral particles and cell lysates and made three key discoveries: (i) FCS cleavage was detected in viral particles across all lineages tested (Fig. 2A), with increasing frequency upon the appearance of Omicron lineages (as assessed with the ratio of cleaved to uncleaved S); (ii) a continuum of virion S protein loss was observed, peaking upon the arrival of XBB.1 and JN.1 Omicron sub-lineages in the VeroE6-ACE2-TMPRSS2 line (Fig. 2A); (iii) infected cell lysates contained a constant pool of uncleaved S protein for all lineages (Fig. 2B). This uncleaved S pool represented an enrichment relative to cleaved forms in Omicron lineages, possibly reflecting the specific depletion of cleaved S in intracellular compartments.

**Figure 2.**
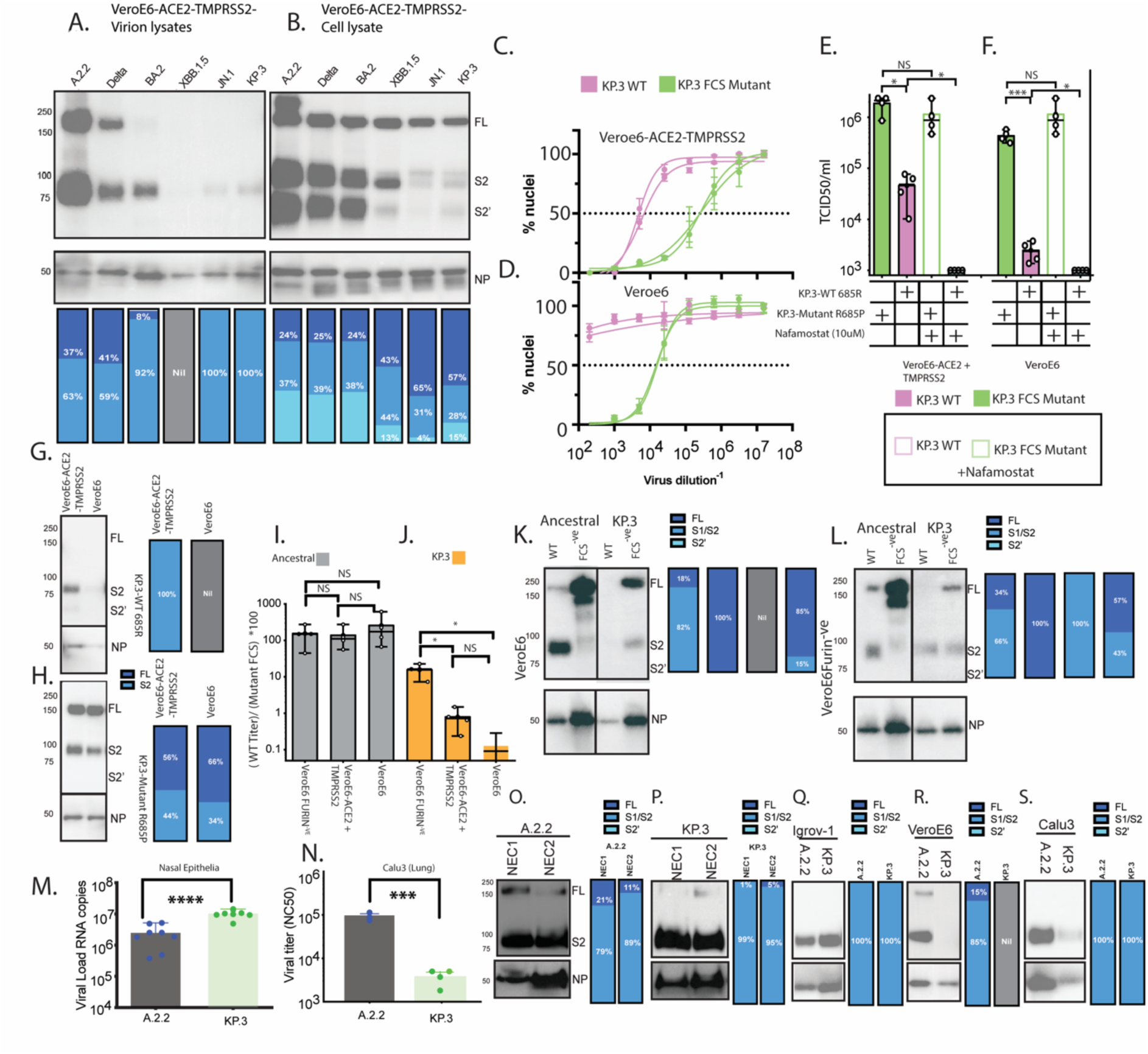
Furin like protease hyper-cleavage drives reduced Omicron S protein virion content in cell type dependent manner and is independent of TMPRSS2. **A. & B.** Representative pre-Omicron and Omicron lineages expanded in the ACE2-VeroE6-TMPRSS2 line for 24 hours at a MOI of 0.025. S & N viral proteins are then analysed by immunoblotting using SARS-CoV-2 Spike Protein S2 antibody (1:2000, MA535946, Thermo Fisher), and anti-SARS-CoV-2 Nucleoprotein (1:2000, Cellabs) from lysates derived in **A.** viral particles or **B.** cell lysates. Data is representative of three independent viral expansions. Molecular weights for proteins are to the left and are in KDa. FL = Full length S protein, S2 = FCS Cleaved and S2’ S2 cleaved. Proportion of each band via densitometry is summarised in blue shaded histograms. **C&D.** Viral titrations with and without the TMPRSS2 inhibitor Nafamostat based on dose-dependent nuclei loss for representative Omicron WT KP.3 (pink) and the FCS resistant mutant KP.3^FCS^ ^mutant^ (green) in **C.** VeroE6-ACE2-TMPRSS2 line and **D.** Unmodified parental VeroE6 line that expresses low endogenous ACE2 but negative for TMPRSS2. Two independent experiments are presented as two independent sigmoidal curve fits. Each point represents the mean with error bars standard deviations of 8 technical replicates. **E.** Summary of Viral titers of WT KP.3 and KP.3^FCS^ ^mutant^ calculated through TCID50/ml on VeroE6-ACE2-TMPRSS2 line & unmodified VeroE6 line with and without Nafamostat. **** P<0.0001; Mann Whitney. Each point represents an independent experiment titer. Error bars are standard deviations across independent experiments. Analysis of **G.** KP.3 WT & **H.** KP.3^FCS^ ^mutant^ viral particles following expansions in **VeroE6-**ACE2-TMPRSS2 VeroE6 and VeroE6 cells as outlined in A. Molecular Weights in KDa right of each panel. Data is representative of three independent viral expansions. Proportions of each band presented as in A&B. **I.&J.** Expansions and relative titers in VeroE6-ACE2-TMPRSS2, VeroE6 and VeroE6 Furin knock out lines (VeroE6FURIN^-ve^) for **I.** KP.3 and **J.** WT Ancestral relative to their FCS mutant controls. Viral supernatant is titered using the equi-permissive VeroE6-TMPRSS2 line. Data is expressed as WT/FCS mutant*100. 100 represents WT equivalence to the matched FCS mutant. Each point represents an independent experiment titer. Error bars are standard deviations across independent experiments. *P<0.05; Student’s unpaired *t-*tests. **K.&L.** Viral expansions of Ancestral, Ancestral^FCS^ ^mutant^, KP.3, KP.3^FCS^ ^mutant.^ in **K.** VeroE6 furin positive or **L.** VeroE6FURIN^-ve^. Viron lysates are presented for K&L with detection of S & N viral proteins as outlined in A&B. Molecular weights to the left in KDa and viral proteins indicted on the right as in A&B. Proportions of each band presented as in A&B. **M.&N.** Contrasting replication of Ancestral vs Omicron KP.3 in **M.** 8 independent donors of nasal epithelial cells versus **N.** the lung derived Calu3 cell line. Each point in M. is an independent donor and in N. an independent experiment with the same cell line. **O.** to **S.** S & N protein products in viral lysates from infections with **O.**&**P.** primary nasal epithelial cell organoids. D1and D2 represent material processed from two independent donors from **M., Q.** Igrov-1 ovarian derived cell line, **R.** Unmodified VeroE6 & **S.** lung derived Calu3. All experiments here are representative of three viral expansions. Molecular weights to the left in O. and in KDa, with viral proteins indicated as per panel B. Molecular weights and viral proteins in O. then also consistent with panels P. through to S. Proportions of each band presented across O. to S. as in A&B.

To test whether S protein FCS cleavage drives virion S protein depletion, we focused on the extremes of this continuum, through comparing the Ancestral lineage with the JN.1 sub-lineage KP.3. Furthermore, to resolve the role of the S FCS site, we obtained both Ancestral and KP.3 S FCS inactivating mutations through 6 rounds of VeroE6 serial passage^24^. For KP.3, FCS inactivation was obtained through acquisition of S R685P and in contrast Ancestral inactivation was obtained by ^679^NSPRRAR^685^ deletion. The KP.3 FCS mutant (KP.3^FCS^ ^mutant^), alongside wild type (WT) KP.3, were then inoculated into both the VeroE6-ACE2-TMPRSS2 line and unmodified VeroE6 cells as controls (Fig. 2C&D). KP.3^FCS^ ^mutant^ rescued viral titers in the VeroE6-ACE2-TMPRSS2 line by approximately 35-fold compared to WT KP.3 (Fig. 2C–E). Parallel analysis in unmodified VeroE6 cells showed a striking phenotype, by which these cells were refractory to WT KP.3 infection but fully permissive to the KP.3^FCS^ ^mutant^ (Fig. 2C-F). Nafamostat treatment demonstrated that KP.3^FCS^ ^mutant^ was resistant to TMPRSS2 inhibition (Fig. 2 E&F). Analysis of virion-associated proteins in Omicron KP.3 revealed partial or complete S depletion across both ACE2-VeroE6-TMPRSS2 and unmodified VeroE6, whereas S amounts were rescued in the KP.3^FCS^ ^mutant^ (Fig. 2 G&H). These results indicate that the FCS is essential for S protein depletion, since inactivation of the FCS completely restored S virion content and infectivity across both cell lines for Omicron KP.3. Of further note, in infected cell lysates across all lineages, there was a constant fraction of S protein that bypassed FCS cleavage and was preferentially enriched in Omicron lineages due to cleaved S depletion.

To determine whether furin, rather than other proteases, was responsible for S protein depletion, we utilised furin-knockout (K/O) VeroE6 cell lines ^7^. In furin-deficient cells, KP.3 was partially rescued, whereas in furin-proficient VeroE6 cells, KP.3 showed significant attenuation (Fig. 2I). In contrast, Ancestral controls observed similar levels of infection across all conditions (Fig. 2J). Omicron KP.3 virion analysis confirmed this proteolytic specificity: in furin-KO cells, S protein was preserved on both KP.3 and KP.3^FCS^ ^mutant^ virions, whereas in furin-positive cells, S protein was selectively depleted from KP.3, but maintained for the KP.3^FCS^ ^mutant^ virions (Fig. 2K&L). In Ancestral controls, there was limited depletion of FCS cleaved S in VeroE6 but decreasing FCS cleaved S in VeroE6-Furin^-ve^ in favour of full-length S (Fig. 2K&L). These findings establish that (i) furin can contribute to protease driven S protein depletion from Omicron virions; (ii) S cleavage of Omicron KP.3 FCS can proceed independently of Furin; and (iii) Ancestral S is relatively resistant to protease S depletion and is observed to have greater dependence on furin for FCS-Cleavage. From the above observations for Omicron KP.3, we hereon refer to this phenotype as protease FCS hyper-cleavage, which is the proteolytic processing of the S protein FCS within cells that culminates in its depletion from viral particles.

To determine whether S protein virion loss was cell-type-dependent, we examined S protein content across cell cultures in which Omicron was either more or less fit than pre-Omicron lineages. Permissive tissues included primary nasal epithelial cells (NEC) from eight independent donors and the ovarian cell line IGROV-1^25^. Low-to-non permissive tissues included VeroE6 and the lung-derived Calu3 cell lines. Infection of NEC organoids confirmed that Omicron KP.3 replicated significantly more efficiently than pre-Omicron Ancestral (Fig. 2M). In contrast, Calu3 cells showed significant KP.3 attenuation (Fig. 2N). Virion analysis revealed striking cell-type specificity: NEC and IGROV-1 virions showed equivalent S protein levels for both Ancestral and KP.3 (Fig. 2O–Q), whereas VeroE6 and Calu3 demonstrated S protein depletion in Omicron KP.3 but not Ancestral virions (Fig. 2 R&S). Critically, this cell-type specificity was independent of TMPRSS2 expression, with two TMPRSS2 negative lines (Human Tissue Atlas ^26^: ENSG00000184012) observing polarised outcomes with respect to virion S protein depletion: VeroE6 cells sustained S protein depletion, whereas IGROV-1 cells were resistant. These results established that protease S FCS hyper-cleavage operated independently of TMPRSS2 and was likely regulated by cell-intrinsic factors that determine FCS accessibility to furin and other related proteases in the trans-Golgi network.

To reconcile why this phenotype was not captured by pseudotyping assays, we examined S protein-pseudotyped VSV particles (PV). In contrast to primary SARS-CoV-2 isolates, VSV pseudotypes bearing Omicron S showed neither S protein depletion nor evidence of impaired FCS cleavage (Fig. S1A–B). This discordance was readily explained by fundamental differences in S protein trafficking and viral assembly. PV bud from the plasma membrane and incorporates a S protein with a significant deletion to its cytoplasmic tail ^27^ and in the absence of the SARS CoV-2 viral Membrane protein. In contrast for authentic SARS-CoV-2, full length S protein enables binding to ezrin-radixin-moesin (ERM) proteins via its cytoplasmic tail and alongside expression of the SARS-CoV-2 Membrane protein, enables an equilibrium of S trafficking to either the membrane of infected cells or SARS-CoV-2 assembly sites ^28^. Given the complexity of the latter, it is consistent divergent pathways of S protein trafficking culminating in differing outcomes to PV and SARS-CoV-2 infections.

Interestingly, replication outcomes in ACE2-TMPRSS2-co-expressing cells were entirely concordant with S protein loss from virions and attenuation of cell-free spread. In contrast, this concordance was disrupted in TMPRSS2-only overexpressing VeroE6 cells (which have low endogenous ACE2) and Omicron lineages that produced S protein-depleted virions were rescued. As the critical difference was ACE2 expression relative to TMPRSS2 levels, we hypothesized that ACE2 and TMPRSS2 dynamics can further influence Omicron replication outcomes alongside the impact of virion S depletion. Thus, we investigated the relationship between ACE2 and TMPRSS2 and how it influenced SARS-CoV-2 infection.

### The ACE2 Collectrin Like Domain (CLD) gates TMPRSS2 usage and rescues Omicron infection *in vitro* and *in vivo*

For initial analysis of ACE2-TMPRSS2 dynamics, we analysed ACE2 shedding, which can be driven by ACE2 cleavage by ADAM17 or other proteases, potentially including TMPRSS2, at the ACE2 collectrin-like domain ^29,30^. As a surrogate of ACE2 shedding, we detected the enzymatic activity of soluble ACE2 in the supernatants of several cell lines and observed lack of soluble ACE2 specifically in the VeroE6-ACE2-TMPRSS2 line (Fig. S2 A–C). In agreement with this finding, there was no evidence for ACE2 cleavage in this cell line (Fig. S2C). In contrast when we transiently expressed ACE2 and TMPRSS2 in HEK293T cells we observed ACE2 cleavage by TMPRSS2 (Fig. S4 A&B), highlighting differential outcomes in this latter cell line as previously described^29^.

Residues in the CLD can regulate TMPRSS2-mediated ACE2 interactions, which has previously been shown to augment SARS-CoV-1 entry into cells ^29^. Given this latter precedent, we next investigated whether mutations in the CLD that abrogated TMPRSS2-mediated ACE2 events (mutant C4 and derivatives ^29^) could impact both ACE2-TMPRSS2 binding and SARS-CoV-2 infection (Fig. 3A-C). ACE2 mutant C4 and its derivatives (Fig. 3B&C) were compatible with TMPRSS2 binding, as determined by immunoprecipitation (Fig. 3D) and biolayer interferometry analyses (Fig. 3E) whereas TMPRSS2 binding to WT ACE2 was markedly reduced (Fig. 3D & E). Mass photometry analyses of these soluble ACE2 ectodomain constructs demonstrated predominant dimer formation of WT ACE2 relative to C4 mutants which were differentially enriched for ACE2 protomers (Fig. 3F) and thus support ACE2 related oligomerisation events can influence TMPRSS binding.

**Figure 3.**
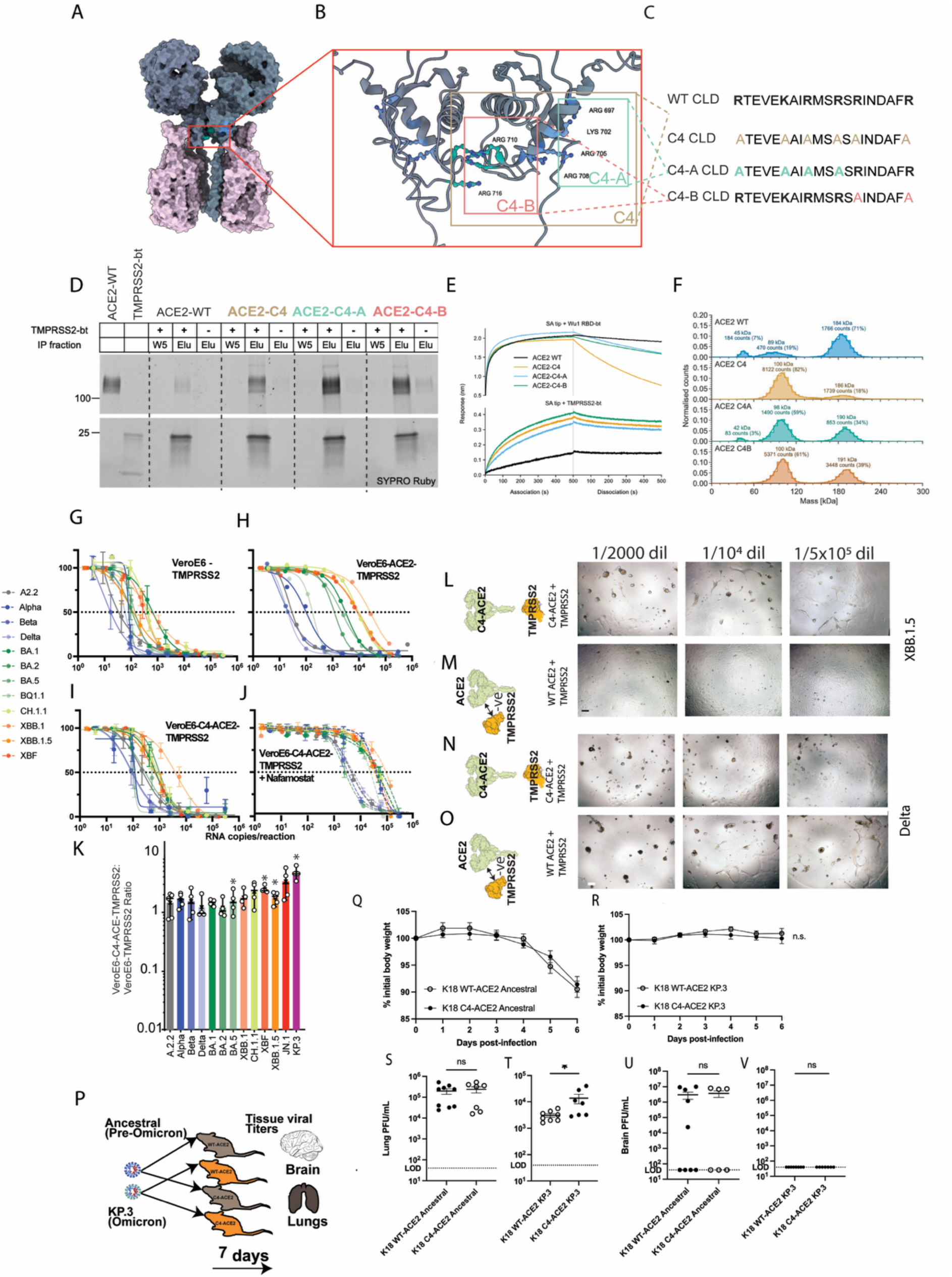
ACE2 CLD mutants promote replication in Omicron lineages. **A.** ACE2 structure with previously proposed TMPRSS2 binding site boxed in red (Structure PDB 6M17). **B**. Expanded view of the CLD domain with C4-ACE2 residues. **C.** CLD-mutants C4, C4-A & C4-B. Alanine substitutions across the CLD are presented in yellow for C4-ACE2, blue for C4-A & orange for C4-B. **D.** TMPRSS2 (S441A) ACE2 pull downs across the listed C4 mutant panel in C. ACE2 runs near 100 kDa in the upper part of the gel. The TMPRSS2 N-terminal fragment runs near 25 kDa, while the biotinylated C-terminal fragment remains immobilized on the resin and is therefore not observed here. ACE2 and TMPRSS2 standards contain 100 ng protein, reduced with 20 mM DTT We note that the amounts of ACE2 pulled-down are small compared to total input. **E.** BLI on the mutant panel presented in C. Upper traces represent ACE2 binding to immobilized Ancestral Wu-Hu-1 RBD. Lower panels represent ACE2 binding to immobilized TMPRSS2 (S441A). **F.** Mass photometry counts for the ACE2 CLD mutant panel presented in D. **G.-J.** Viral titration curves as initially outlined in Fig. 1B&C, across 12 primary SARS-CoV-2 isolates with **G.** VeroE6-TMPRSS2, **H.** VeroE6-ACE2-TMPRSS2, **I.** VeroE6-C4-ACE2-TMPRSS2 **J.** VeroE6-C4-ACE2-TMPRSS2 +10μM Nafamostat. Input viral inoculum in G.-J. is presented here in RNA copies per infection. Sigmoidal dose-dependent curves represent cell loss from viral cytopathic effects (% nuclei). Standard deviations in each curve are derived from 8 technical replicates. Titration curves presented are representative of three independent experiments. **K.-N.** Contrasting replication across variants appearing in WT ACE2-TMPRSS2 and C4-ACE2-TMPRSS2 cell lines presented, for Omicron lineage XBB.1.5 (K.&L.) and the pre-Omicron lineage Delta (M.&N.). Images depict viral titrations at three select viral dilutions and are representative of greater than 3 independent experiments. Scale bar: 20 µM. **O.-T.** Two transgenic lines were established to validate *in vitro* observations *in vivo* and include **O.** K18-WT ACE2 and C4-ACE2 mice. Both lines were infected with either Ancestral (Clade A.2.2) or Omicron lineages (KP.3). **P.&Q.** Weights were then monitored for 7 days post infection after which **R.&S.** Lung and **T.&U.** Brain were harvested to titer viral loads in each tissue. Weight curves were analysed using a two-way ANOVA, while viral titers were analysed using Student’s unpaired *t-*tests.

Although the dynamics of ACE2-TMPRSS2 interactions extend beyond initial binding alone, with the CLD known to contribute to augmented SARS-CoV-1 entry^29^. As with SARS-CoV-1, we tested if the CLD differentially affects SARS-CoV-2 infection primarily through the use of C4-ACE2 mutants. Omicron sub-lineages infected C4-ACE2/TMPRSS2 cells more efficiently than WT ACE2/TMPRSS2 cells and infectivity was Nafamostat sensitive, confirming that entry was TMPRSS2-dependent (Fig. 3G–O). To complement soluble ACE2-TMPRSS2 studies, we turned to analysis of potential ACE2-TMPRSS2 complexes in live cells. In contrast to soluble protein analysis, expression of WT ACE2 led to detectable ACE2 in TMPRSS2 pulldowns, but this was subsequently reduced in C4-ACE2 mutants (Fig. S4. C-E). As ACE2 on live cells exists as a mixture of monomeric and oligomeric species^31^, the combined observations of soluble proteins versus transient expression in live cells supports TMPRSS2 engagement with primarily monomeric ACE2, with residues with the CLD further influencing ACE2-TMPRSS2 complexes. The functional effects of ACE2 CLD mutants thus likely run through modulation of TMPRSS2 binding (or processing), whether directly or indirectly through altered dimer equilibria. Given the results *in vitro* with C4-ACE2 mutants, we then validated in lung tissue *in vivo* by the generation of two transgenic mouse lines that expressed either WT ACE2 or C4-ACE2 (Fig. 3P). In this setting, the ACE2 CLD is highly conserved among human and mouse ACE2 and human ACE2 is processed by endogenous murine proteases of the renin-angiotensin system ^32^. Each transgenic line was infected with either Clade A.2.2/Ancestral or the Omicron sub lineage KP.3. Ancestral virus replicated in both K18-WT ACE2 and K18-C4-ACE2 mice to comparable titers and induced weight loss with no difference between groups (Fig. 3Q). In contrast, infection with KP.3 did not result in weight loss across both murine lines (Fig. 3R), as expected given the attenuation of Omicron sub lineages. Notably, lung viral loads were significantly higher in the C4-ACE2 versus WT ACE2 murine line (Fig. 3S&T), but not to the levels observed with the Ancestral Clade A.2.2. Brain viral loads were only detected in Ancestral Clade A.2.2 infections (Fig. 3U&V). These findings support two observations: first, heavily attenuated KP.3 Omicron replication in the lung was consistent with outcomes in the lung-derived human Calu3 cell line and S protein virion depletion; second, mutations modulating ACE2-TMPRSS2 interaction can rescue viral infection *in vitro* and *in vivo*.

### ACE2-associated solute carriers rescue Omicron sub-lineage infection in a TMPRSS2 dependent manner

We then investigated orthogonal physiological conditions that may also regulate TMPRSS2 access to ACE2. ACE2 can act as a chaperone for the tissue-specific solute carriers SLC6A19^33,34^ and SLC6A20 ^35–37^ which bind to the ACE2 TM domain and CLD ^36,38–40^ and thus possibly interfere with TMPRSS2 binding, again either directly or indirectly through ACE2 dimerization. To investigate the potential for solute carriers to influence SARS-CoV-2 infection in relevant permissive tissues, single cell level analysis was performed and revealed the co-expression of ACE2, TMPRSS2 and SLC6A19/SLC6A20 across tissues known to be targets for SARS-CoV-2 infection: ACE2^++^/SLC6A19^++^/TMPRSS2^++^ intestinal enterocytes, ACE2^+/-^/SLC6A20^+/-^/TMPRSS2^++^ lung type 2 pneumocytes, and ACE2^+^/ SLC6A20^++^/TMPRSS2^++^ ciliated nasal epithelia (Fig. S5).

We next investigated whether solute carriers impact SARS-CoV-2 infection. Since overexpression of solute carriers relative to ACE2 can lead to ACE2 cytosolic retention ^36^, we ensured co-expression of ACE2 and solute carriers at comparable levels in the clonal VeroE6 cell lines TASL-19 (TMPRSS2+ACE2+SLC6A19) and TASL-20 (TMPRSS2+ACE2+SLC6A20) (Fig. S6 A–D). Both pre-Omicron lineages and Omicron sub-lineages replicated in the TASL-19 cell line with higher efficiency as compared to the parental VeroE6-TMPRSS2 cell line (Fig. 4 A&B). Similarly, Omicron sub-lineages circulating in late 2025 could be isolated with high and comparable efficiency using TASL-19 cells (Fig. S6E). Finally, infection of TASL-19 cells was inhibited by Nafamostat across all pre-Omicron and Omicron lineages with similar potency (Fig. 4C). Collectively, we found that co-expression of SLC6A19 with ACE2 and TMPRSS2 allowed for robust and comparable infection with pre-Omicron and Omicron sub-lineages in a TMPRSS2-dependent manner.

**Figure 4.**
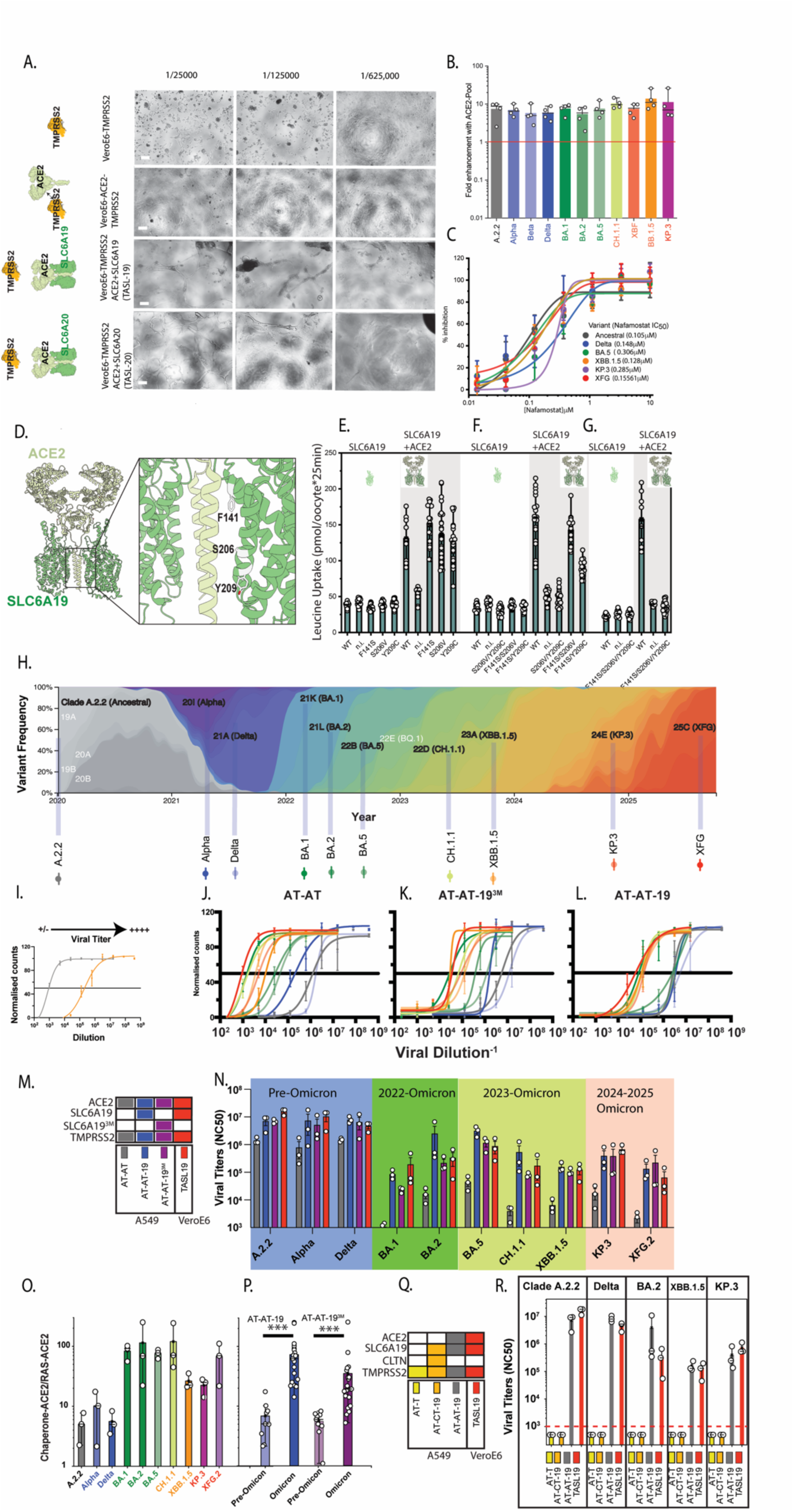
Solute carriers chaperoned by ACE2 augment infection across all SARS-CoV-2 lineages. **A.** Cells were inoculated with the KP.3 Omicron isolate at dilutions stated above panels. Within 48 hours, extensive syncytia is observed primarily in TASL-19 and TASL-20 cell lines. Limited to no cytopathic effects are observed using the VeroE6-ACE2-TMPRSS2 cell line at equivalent viral inocula. Images are representative of three independent experiments. Scale bars are at 50μm. **B.** Summary of fold augmentation of all major variants to date of the TASL-19 cell line. Fold augmentation is calculated as the fold change of the engineered TASL-19 line compared to the parental cell line VeroE6-TMPRSS2. Data points are presented from independent experiments. **C.** TMPRSS2 dependence in TASL-19 across representative pre-Omicron and Omicron lineages. Nafamostat dose-dependent inhibition of the TASL-19 line as previously described ^3^. Interpolated [Nafamostat] that sustain 50% inhibition are listed alongside each isolate. Each data point represents four technical replicates with standard deviations as error bars. Data presented are representative of three independent experiments. **D.**-**G.** SLC6A19 mutants and their ability to sustain amino acid transport. **D.** Mutants that sustained ACE2 binding for their ability to transport amino acids. **E.** Amino acid transport in single mutants, **F.** double mutants and **G.** Triple mutants. **I.**-**N.** Triple SLC6A19 mutants were then selected to observe their capacity to enhance viral infection across a panel of variants that span the pandemic. **H.** The A549 lung-derived cell line was selected to test combinations of ACE2, SLC6A19 and TMPRSS2, as they do not express these factors endogenously. In each case, cells are transduced using lentiviral vectors, clonally sorted and expanded and selected based on expression of each factor (Fig. S5). **I.** Viral titers were screened as outlined in Fig. 1. Here, dose-dependent loss of nuclei is used to quantify viral replication following a 48 hour culture. **J.** Replication of variants with the AT-AT cell clone: A549 with ACE2+TMPRSS2 expression (RAS-ACE2). **K.** The AT-AT-19 clone: ACE2+SLC6A19 +TMPRSS2 **L.** The AT-AT-19^3M^ clone: ACE2+SLC6A19 triple mutant +TMPRSS2. In L.-N. standard deviations represent 4 technical replicates. **M.** Expression of ACE2, solute carrier and TMPRSS2 across each engineered A549 cell line presented in N. to P. **N-P.** From J. to L. three independent experiments are summarised through calculation of viral dilutions that sustain 50% cell loss as estimated by the sigmoidal curve fits. The VeroE6 TASL-19 based cell line is presented here as a positive control to the A549 derived clonal cell lines. **O.&P.** Fold-augmentation of virus replication of Chaperone-ACE2 (ACE2+SLC6A19+TMPRSS2) versus RAS-ACE2 (ACE2+TMPRSS2) is presented from data in **N.** *** p<0.0001 for statistically greater enhancement in Omicron vs Pre-Omicron; Mann-Whitney. For both Chaperone-ACE2 (AT-AT-19) and Chaperone-ACE2 with triple SLC6A19 mutant (AT-AT-19^3M^). **Q&R.** Control A549 clonal lines are generated as in J. to L. but without ACE2 expression. AT-T: TMPRSS2 A549 clone; AT-CT-19: SLC6A19 + collectrin (alternate chaperone)+TMPRSS2 clone. Red dashed line represents the limit of detection. Positive control cell lines AT-AT-19 and TASL-19 were tested in parallel. Each point represents titers from independent experiments.

To resolve if the amino acid transport function of SLC6A19 was needed for supporting efficient Omicron infection, we screened for SLC6A19 mutations that maintain ACE2 binding but are impaired in amino acid transport (Fig. 4D–G). A combination of three mutations within SLC6A19 (F141S/S206V/Y209C; SLC6A193M) stringently ablated amino acid transport (Fig. 4G). To test whether this mutant still promotes SARS-CoV-2 infection, we generated stable A549 lines since these cells lack ACE2, TMPRSS2 and solute carrier expression and were derived from the human lung (Fig. S7). In A549 cells, co-expression of ACE2 and TMPRSS2 allowed for efficient infection with pre-Omicron lineages but not Omicron sub-lineages and thus validated our findings made with VeroE6 cells. Co-expression of ACE2 and SLC6A19 jointly with TMPRSS2 augmented infection across all SARS-CoV-2 lineages approaching levels observed in the VeroE6-based TASL-19 cell line (Fig. 4I–P). Expression of SLC6A19^3M^ with ablated amino acid transport capacity also allowed for efficient infection at levels similar to those measured for A549 cells expressing WT SLC6A19 (Fig. 4N). Finally, we tested if solute carriers alone could support infection when co-expressed with TMPRSS2. For this, we substituted ACE2’s chaperone function by co-expression of Collectrin at equimolar ratios with SLC6A19. When ACE2 was not expressed (TMPRSS2 alone and Collectrin+SLC6A19+TMPRSS2) no infection was detected (Fig. 4Q&R). These findings are important at two levels. Firstly, infection across all SARS-CoV-2 lineages remains dependent on ACE2 and TMPRSS2 but is augmented in the presence of solute carriers. Secondly, amino acid transport activity of SLC6A19 is dispensable during this viral augmentation.

### Free TMPRSS2 cleaves the S protein FCS and promotes SARS-CoV-2 replication during cell-cell spread

We next addressed the underlying mechanisms of ACE2-related solute carrier augmentation. To investigate this, we analyzed outcomes across key cell lines using standard plaque assays. Plaque assays comparing lines with and without solute carriers revealed significantly larger plaques across all lineages for the SLC6A19 positive TASL-19 cell line (Fig. 5. A–C). In contrast Omicron lineages were significantly attenuated in both titer and plaque size in the VeroE6-ACE2-TMPRSS2 line. Given S protein depletion in VeroE6 derived cells is observed in Omicron virions, the plaque size likely reflects the efficiency of cell-cell spread. To resolve this further, we investigated cell-cell spread efficiency in our cell line model systems, focusing on S FCS cleavage of the S protein in cell lysates as opposed to viral particles.

**Figure 5.**
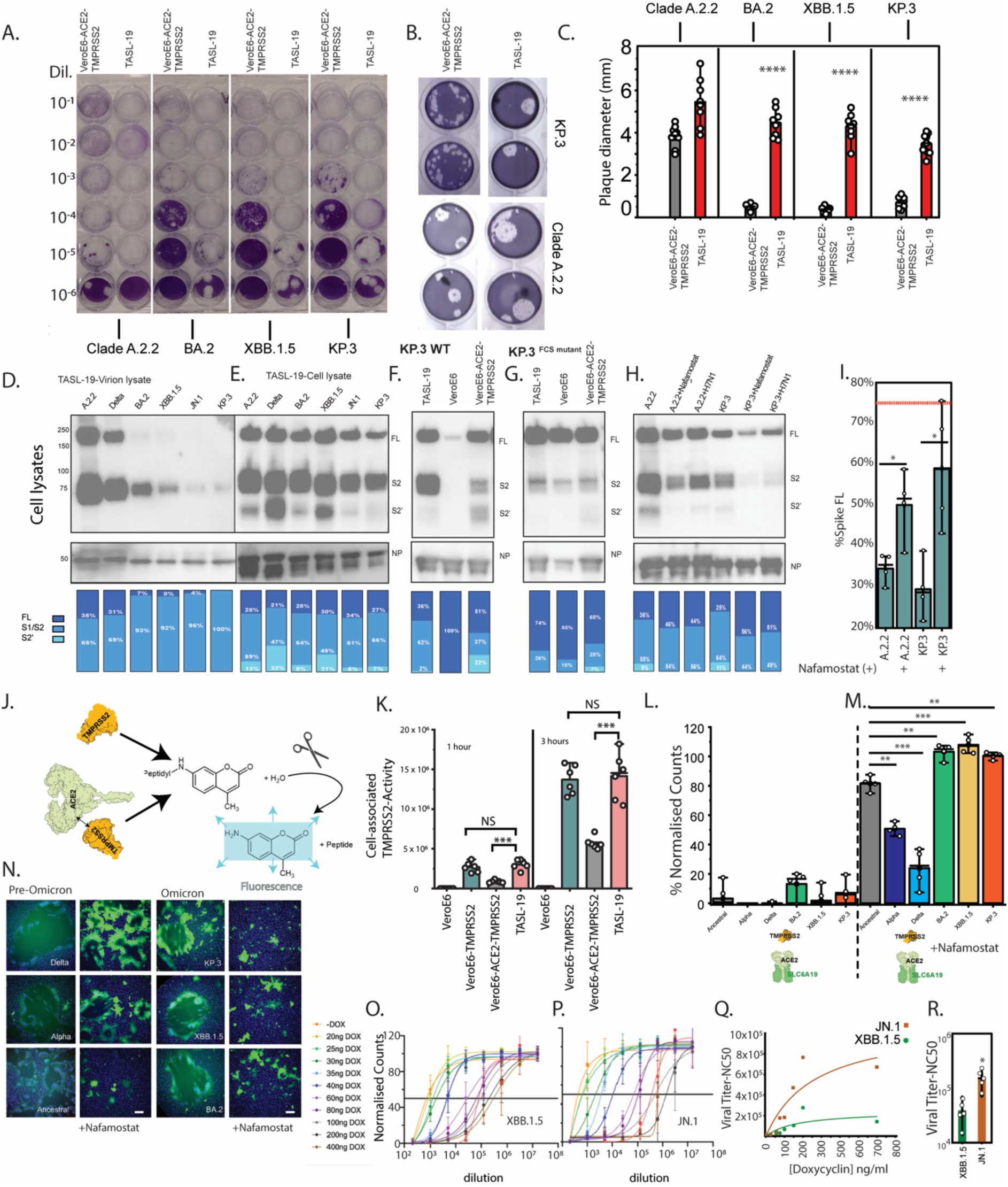
TMPRSS2 recognises the S protein FCS and is primarily used by Omicron lineages for cleavage during cell-cell spread. **A.** Viral plaque assay for representative pre-Omicron and Omicron lineages in VeroE6-ACE2-TMPRSS2 and TASL-19. Viral dilution is presented to the left. **B.** Expanded view of viral plaques from an independent experiment with Ancestral Clade A.2.2 and Omicron KP.3. **C.** Plaque diameter calculated from three independent experiments. Each point represents an individual plaque. **D&E.** SARS-CoV-2 listed isolates were inoculated on the **D.** VeroE6-ACE2-TMPRSS2 line or **E.** the TASL-19 line for 1 hour at an MOI of 0.025. 24 hours post-infection cells were lysed and lysates were subject to western blotting for S and N proteins. Data is representative of minimum of 4 separate pre-Omicron and Omicron expansions. Molecular weights are presented on the left of panel I. and are in KDa. Densitometry captures proportion of individual cleavage bands against total bands using ImageJ. Dark blue represents full length spike (FL), light blue represents S1/S2, and cyan represents S2. **F. & G.** Accumulated S and N proteins for KP.3 in TASL-19, VeroE6 and VeroE6-ACE2-TMPRSS2 lines under conditions outlined in D&E. with the **F.** S protein FCS intact or **G.** S protein FCS with the R685P mutation. **H.** Expansion of virus with the anti-TMPRSS2 antibody (H171) and Nafamostat added post-entry for Ancestral Clade A.2.2 and KP.3. Data is representative of three separate viral expansions from **D. to H.** I. Summary of full-length Spike with and without the addition of Nafamostat post-entry. Each point represents an independent viral expansions in the TASL-19 cell line. Red line represents the proportion of full-length S protein for the KP.3 R685P mutant * P<0.01, Mann-Whitney. **J.&K.** Cell associated TMPRSS2 measured through cell culture of the **J.** Boc-Gln-Ala-Arg-AMC fluorogenic peptide for **K.** TMPRSS2 negative VeroE6, VeroE6-TMPRSS2, VeroE6-ACE2-TMPRSS2 and the TASL-19 cell line. Each data point represents independent experiments with error bars standard deviation from the mean. *** p<0.0001; Mann-Whitney. **L.&M.** Transfection of S protein and eGFP in the TASL-19 line reveals cell-cell fusion **L.** without and **M.** with the TMPRSS2 inhibitor Nafamostat through cell nuclei loss. **N.** Representative images from transfections in **L.&M.** Scale bars are at 50uM. **O-Q.** Establishment of the Tet-TASL-19 cell line with regulation of TMPRSS2 levels alongside Chaperone ACE2 (ACE2+SLC6A19). Viral titers presented as dose dependent nuclei loss for each Doxycycline (DOX) condition for **O.** Omicron XBB.1.5 and **P.** Omicron JN.1. **Q.** Interpolated 50% nuclei loss values are calculated from sigmoidal curves presented in O. and P. **R.** 5 independent titers established with 100ng/ml Doxycycline for XBB.1.5 and JN.1. *<0.01; T-Test.

As with the VeroE6-ACE2-TMPRSS2, the TASL-19 line generated Omicron viral particles subject to hyper-cleavage and S protein depletion (Fig. 5D). However, in cell lysates, FCS-cleaved S was substantially rescued and equivalent across all lineages (Fig. 5E). Furthermore, in cell lysates the KP.3^FCS^ ^mutant^ abrogated S FCS processing (Fig. 5F & G). These findings support the concept that S protein content and FCS processing can be rescued upon co-expression of ACE2 and SLC6A19 with TMPRSS2, but at the infected cell as opposed to the virion membrane. Alongside plaque assays, this further supports viral rescue of Omicron lineages is primarily during cell-cell spread events. Throughout, we have observed a pool of S that is resistant to depletion in infected cell lysates. Differential trafficking and regulation of this S pool is consistent with surface Son infected cell membranes directing replication outcomes relevant to cell-cell spread but independent of cell-free viral particle spread. TMPRSS2 sustains extracellular activity at the plasma membrane ^41^ and *in vitro* assays with recombinant TMPRSS2 and S provide direct evidence that TMPRSS2 can bind and cleave the S FCS ^42^.

To resolve this further, we blocked TMPRSS2 after viral entry to capture cell-cell spread dynamics. Examination of S FCS cleavage in infected cells treated with either the TMPRSS2 inhibitor Nafamostat or the TMPRSS2 antibody H1H7 post-entry revealed reduced FCS cleavage of Ancestral S and KP.3 S approaching that of the KP.3 FCS mutant (Fig. 5.H&I). This observation is important for two reasons: Firstly, it provides evidence that TMPRSS2 can engage the FCS at the cell surface when solute carriers have engaged ACE2. Secondly, this TMPRSS2-mediated FCS cleavage is gated by ACE2-TMPRSS2 co-expression, supporting ACE2 as a negative regulator of TMPRSS2 FCS cleavage. To test the latter, we measured cleavage of a cell-impermeable fluorogenic TMPRSS2 substrate as a surrogate for surface TMPRSS2 activity (Fig. 5. J). Whilst surface TMPRSS2 activity was readily detected in VeroE6-TMPRSS2 and TASL-19 cells, it was significantly reduced in VeroE6-ACE2-TMPRSS2 cells. This directly supports the ability of ACE2 to negatively regulate TMPRSS2 activity on cleavage substrates at the cell surface (Fig. 5. K) and further supports the hypothesis that ACE2 can gate S FCS cleavage by TMPRSS2. In contrast the entry of equivalent variant pseudotypes remains unimpeded in the VeroE6-ACE2-TMPRSS2 line and thus supports the gating of TMPRSS2 by ACE2 doesn’t extend to negative regulation of S2 cleavage.

To further investigate the role of TMPRSS2 activity towards S at the cell surface, we focused on S protein dynamics by expressing S proteins in isolation in the TASL-19 line and then measuring S protein-driven cell-cell fusion in the presence or absence of the TMPRS2 inhibitor Nafamostat. Across all pre-Omicron and Omicron lineages extensive cell-cell fusion was detected. Omicron S protein-mediated fusion was highly dependent on TMPRSS2 activity (Fig. 5. L-P), while pre-Omicron S proteins could sustain cell-cell fusion in a partially TMPRSS2-independent manner (Fig. 5. L-N). This supports that Omicron lineages have evolved to proteolytically restrict S FCS cleavage at the cell membrane to free active TMPRSS2.

As the rescue of Omicron lineages is primarily through surface active TMPRSS2, it was important to establish TMPRSS2 use by Omicron was efficient and not a consequence of TMPRSS2 over-expression. To establish this control ACE2+SLC6A19+ was initially co-expressed in the VeroE6 line to limit ACE2 regulation of TMPRSS2. We then subsequently incorporated a doxycycline-regulated TMPRSS2 construct to generate the clonal cell line Tet-TASL-19 (Fig. S8. A–D). Omicrons XBB.1.5 and JN.1 were then selected based on extensive Spike virion depletion to ensure TMPRSS2 use during cell-cell spread would be observed. Doxycycline tuning of TMPRSS2 levels in the Tet-TASL-19 line led to a dose-dependent effect on Omicron replication, where ∼3-fold reduction in TMPRSS2 surface levels corresponded to a ∼20–26-fold decrease in viral titers (Fig. 5O-Q), consistent with a high dependence of TMPRSS2 use by Omicrons XBB.1.5 and JN.1. Interestingly, we observed JN.1 to utilize TMPRSS2 more efficiently than XBB.1.5 (Fig. 5Q&R).

### Rapid resolution of Omicron phenotypes using clinical material

As outlined above, two overlaid spike protein mechanisms—FCS-driven S depletion and TMPRSS2-dependent cell-cell spread—generate complex but distinguishable outcomes across cell culture systems. While these patterns are evident in in vitro-expanded clinical isolates (attenuation of Omicron lineages in the VeroE6-ACE2-TMPRSS2 line but significant rescue in the TASL-19 line), our aim was to translate this into a rapid clinical assay using patient nasopharyngeal swabs obtained during diagnostic testing and genomic surveillance.

Initially we validated the approach using pre-Omicron lineages and observed no significant difference in titers across both cell lines with Delta swabs (Fig. 6A.; *p* = 0.932). To apply this now to Omicron lineages, we titered swabs alongside COVID-19 genomic surveillance activities (n = 618; surveillance period: April 2025–June 2026) across VeroE6-ACE2-TMPRSS2 and TASL-19 cell lines. During this period, isolates were dominated by Omicron sub-lineages, primarily JN.1 and the divergent BA.3.2.2 lineage (Fig. 6B.). In contrast the Delta swabs, aggregate data form Omicron lineages revealed two orders of magnitude reduction in viral titer in VeroE6-ACE2-TMPRSS2 relative to TASL-19 (*p*<0.0001), consistent with results from in vitro-expanded clinical isolates (Fig. 6C.&D.). Importantly, this distils the *in vitro* complexity of the Omicron phenotype into a 48-hour readout using material already procured for diagnostic testing and genomic surveillance. Additionally, this approach combined with genomic surveillance can facilitate rapid detection of potential phenotypic reversion to the pre-Omicron phenotype, which then can be importantly link linked to changes in clinical outcomes. In such cases, escalation to variant analysis in primary organoid and/or animal models would then proceed alongside clinical observations to confirm any significant tropic shifts.

**Figure 6.**
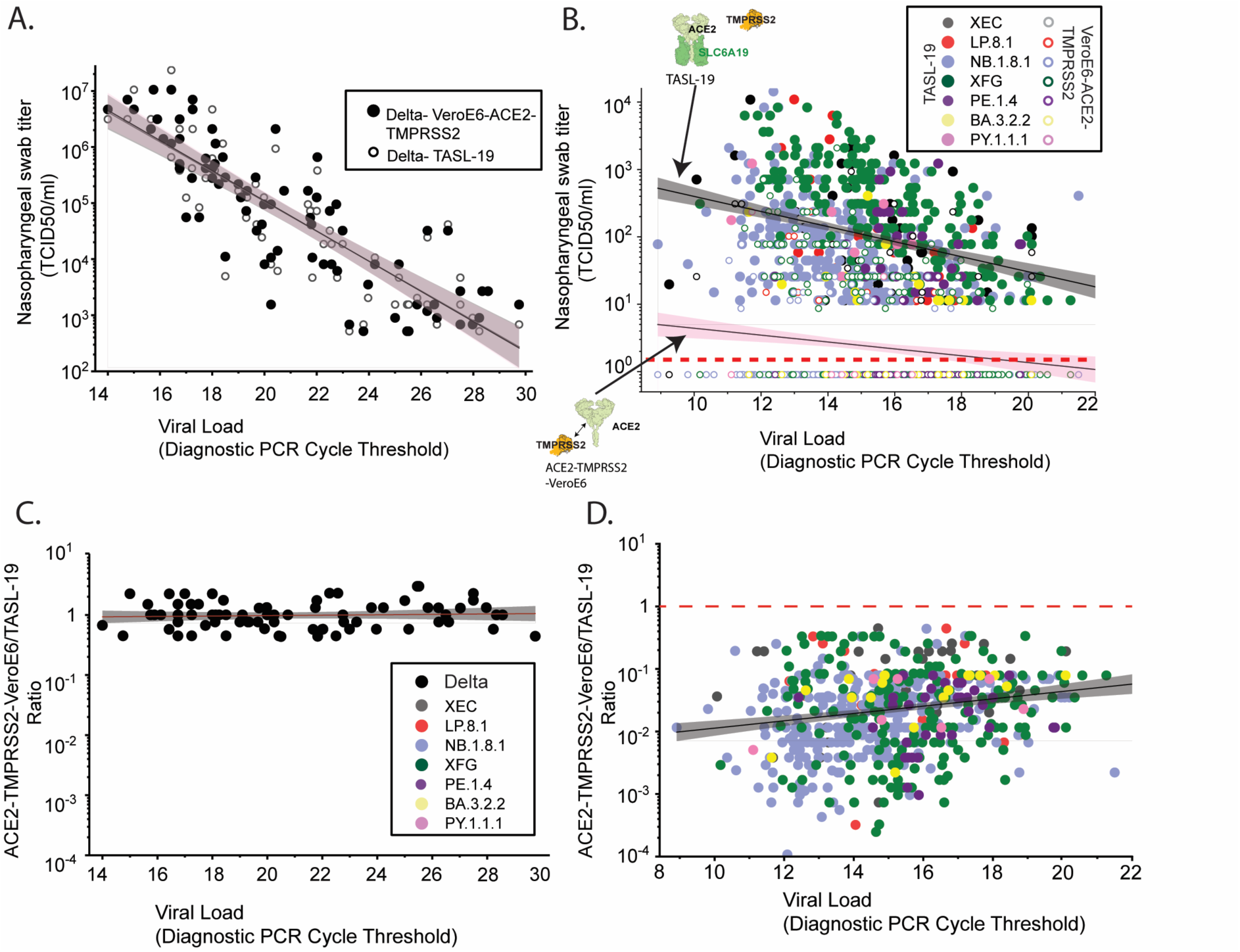
Translation of mechanistic observations herein for rapid resolution of Omicron phenotypes in clinical samples. **A. & B.** Clinical material is derived from nasopharygeal swabs which are initially acquired for respiratory viral testing through diagnostic PCR. SARS-CoV-2 positive samples are then cryopreserved at 80^0^C with a 24-48 hour period, de-identified but with viral load linked to each sample (Diagnostic PCR Cycle Threshold). Small volumes (200ul) of the remnant sample are then processed and titered across the two listed cell lines in tandem (closed circles = TASL-19; Open circles = VeroE6-ACE2-TMPRSS2 in a 384 well plate in quadruplicate. 78 hours post infection, cytopathic effects are visually scored and TCID50 calculated using the Karber method. **A.** Pre-Omicron swabs were obtained in July 2021 when Delta was circulating (n = 72) **B.** For Omicron lineages acquired over the period of April 2025–June 2026 (n = 618), variants are resolved through whole genome sequencing and assigned to nearest parent JN.1 sub-lineage (with the exception of BA.3.2.2). For A. & B. Infectivity per viral load is presented with linear regression fits for all variants pooled displayed with 95% confidence intervals: Grey shaded area = TASL-19 and red shaded area = VeroE6-ACE2-TMPRSS2. Dotted red line represents the limit of detection and points below that line are culture negative but assigned a value of 9. **C. & D.** Data derived in A.&B. is presented as the titer ratio of VeroE6-ACE2-TMPRSS2: TASL-19 versus the diagnostic viral load. In D. The red dashed line signifies the titer ratios Delta swabs in C. Linear fit for all variants pooled are presented with the shaded region representing 95% confidence intervals.

## Discussion

The arrival of Omicron lineages in late 2021 was a seismic phenotypic shift for SARS-CoV-2 and the trajectory of the pandemic^16,17,43^. The preliminary Omicron phenotype proposed at the time was S FCS cleavage resistance and lost pathway flexibility as protease use shifted from TMPRSS2 at the plasma membrane to Cathepsins within endosomal compartments. Whilst the hypothesis was initially supported by *in vitro* observations ^43,44^, additional observations amid evolution of the Omicron phenotype and the Omicron S FCS supported an alternative mechanism that centres around an active FCS in contrast to cleavage resistance, with restored entry pathway flexibility. Here, we observe that outcomes of FCS cleavage changes have driven Omicron’s shift initially with BA.1/BA.2 and then evolved further during the era of Omicron XBB.1 and JN.1 sub-lineages. This optimisation of the Omicron FCS culminates in three outcomes. Firstly, FCS cleavage is optimised at the primary transmission site. This results in virions with maximally cleaved S in the upper respiratory tract and provides the high transmission potential of Omicron lineages. Secondly, the FCS is further consolidated to focus activity towards TMPRSS2-dependent cell-cell spread. Whilst all SARS-CoV-2 lineages benefit from this pathway, Omicron’s further benefit comes from exclusively using it over other proteases. Both pathways then synergise, as one drives the FCS to sustain increased viral particle infectivity, whilst the second ensures cell-cell spread rapidly increases viral load. Importantly, the cell-cell spread pathway is governed by tissue-specific ACE2 ultra-structural organization associated with solute carriers SLC6A20 in the respiratory tract and SLC6A19 in the intestinal enterocytes. Notably, we observe contemporary JN.1 sub-lineages use the latter TMPRSS2 pathway more efficiently during cell-cell spread, and this alone may represent a previously unknown and dominant fitness advantage that coincided with its ability to rapidly supplant co-circulating XBB.1 sub-lineages^45,46^. Finally, the gains in replication fitness in the upper respiratory tract come with opportunity costs elsewhere. Here changes to the FCS make it vulnerable to virion S depletion in cells which subject the FCS to hyper-cleavage. Furthermore, Omicron FCS changes that restrict the use of TMPRSS2 during cell-cell spread and can promote a second layer of attenuation where ACE2-TMPRSS2 complexes form as part of regulating the renin angiotensin system. Whilst this trajectory has enabled the continued emergence of highly transmissible Omicron lineages, consolidation of attenuation in tissues such as the lung has further shaped a trajectory towards lower disease severity.

For S protein regulation on viral particles, in initial studies using isolated Omicron S FCS peptide substrates, maximal furin cleavage was observed^47^, which in isolation is consistent with increased virion infectivity. Yet in situ, Omicron-specific S changes (both directly or indirectly associated with the FCS) manifest in S protein outcomes at several levels. Firstly, the Omicron-specific change at N679K restores S O-glycosylation at position 678^48^ adjacent to the FCS. Secondly, S FCS cleavage outcomes can be influenced by sialylation of carbohydrates at position 678 ^49^. Finally, S protein changes outside of the S FCS that influence S conformational changes may further influence S FCS access to proteases like Furin. The latter is consistent when directly comparing two variants with the same S FCS, BA.2 versus XBB.1, but with very different outcomes related to S FCS cleavage yet with changes primarily in the N-terminal S domain (i.e. XBB>>BA.2 for S depletion from virions). The culmination of these many variables leads to what we presently refer to as the “Goldilocks” FCS model, where FCS cleavage on virions is sustained at an optimised point at the primary transmission site (upper respiratory tract) but comes at the cost of hyper-cleavage at other tissue sites (lower respiratory tract). Cells of the upper respiratory tract, including nasal epithelial cells, are characterized by abundant and heavily sialylated mucin-type O-glycosylation ^50^, which may shield S and lower the intrinsic activity of furin or related proteases towards the Omicron FCS. In contrast, lung epithelial cells, with a distinct glycosylation profile, may have exposed the FCS and made it susceptible to protease hyper-cleavage, culminating in virion S depletion. A simple model for early circulating Omicron lineages predicts a direct correspondence between T678 O-glycosylation status across respiratory tissues and the degree of S depletion observed in virions. Biologically this is consistent with presence of O-GalNAc glycans adjacent to proteolytic cleavage sites regulating the processing of secreted proteins ^51^. Whilst this may account for early Omicron lineages, the consolidation of the Omicron phenotype in Omicron eras dominated by XBB.1 and JN.1 sub-lineages likely reflects the sum of many S changes. For instance, the influence of glycosylation may also extend to cell type specific sialylation of N-glycans on S as revealed in the Omicron specific GWAS signal for ST6 beta-galactoside alpha-2,6-sialyltransferase 1 (SIAT1)^52^. Resolution of the post-translational modifications that influence FCS cleavage represents an important area for future investigation. As is the characterisation of all furin-like proteases that can impact outcomes driven by the S FCS.

For S protein regulation at the infected cell membrane, the outcomes are equally complex. Whilst changes at S position 681 for Alpha and Delta enabled enhanced virion FCS cleavage, they culminated in promiscuous cell-cell fusion across diverse cell types, often independent of TMPRSS2 ^9^. Omicron diverges fundamentally: it combines increased virion FCS cleavage with a restrictive proteolytic strategy where both S FCS and S2 cleavage at the cell membrane are directed exclusively to TMPRSS2. This selectivity—prioritizing TMPRSS2 whilst excluding proteases with sub-optimal outcomes—is adaptive for two reasons. First, the FCS is a finite, consumable substrate; once cleaved, that event cannot be repeated, making it essential to direct it to the most efficient protease. Second, availability to alternative proteases can impose either sub-optimal outcomes or in the worst-case direct S inactivation^15^. For instance, premature cleavage of the S FCS by other proteases may jointly destabilize S and also expose neutralizing epitopes^53^ prior to final engagement of S2 by TMPRSS2. Or alternatively, recognition by other proteases may results in S inactivation (e.g. As observed with HAT^15^). By consolidating cleavage to TMPRSS2, Omicron achieves synchronized S activation at the cell membrane, ensuring that FCS exposure during ACE2 engagement immediately precedes fusion—a temporal coupling that would drive efficient cell-cell spread. Furthermore, continued evolution for optimised targeting of this TMPRSS2-mediated cell-cell pathway may have conferred a transmission fitness advantage on JN.1. Efficient cell-cell spread of virus is a key mechanism used by many viruses to sustain viral replication at orders of magnitude greater than cell-free virus^54^. Furthermore, cell-cell viral transfer can sustain onward transmission for highly contagious viruses like Measles ^55^ and can also enable evasion of neutralizing antibody responses ^56,57^.

However, this TMPRSS2-dependent strategy critically depends on one condition: ACE2 sequestration from TMPRSS2 through solute carrier binding. Whilst the ACE2-solute carrier chaperone complex may compete with TMPRSS2-ACE2 binding, higher-order ACE2 oligomers and ACE2 glycosylation ^58^ may also impact the capacity for TMPRSS2 to bind. Without the presence of solute carriers, ACE2 can bind TMPRSS2 and impede the ability of TMPRSS2 to engage the S FCS. ACE2-solute carrier complexes can exist in the respiratory tract as ACE2-SLC6A20 or alternatively in the intestinal enterocytes as ACE2-SLC6A19. Understanding the opposing outcomes in the upper versus the lower respiratory tract can be explained at several levels. Firstly, the outcomes from two large GWAS studies clearly identify SLC6A20 as a factor that increases transmission across both pre-Omicron and Omicron lineages. Whilst this supports upper respiratory tract infection, augmented infection also within the lower respiratory tract should lead to increased high disease severity outcomes^59^ and this is not observed in pre-Omicron SLC6A20 GWAS observations^60^. Secondly, herein quarantining ACE2 from TMPRSS2 through expression of the C4-ACE2 mutant in mice can rescue lung infection primarily in Omicron lineages. The latter observation importantly highlights that ACE2 has the potential to negatively regulate TMPRSS2 in the lung and as such consolidates Omicron attenuation alongside S protein depletion. Finally, the more efficient use of TMPRSS2 by JN.1 during cell-cell spread manifested in significant dominance over XBB sub-lineages^46^ and was associated with increased transmission and viral loads in faecal swabs^61^ but not linked to increased presentation of high disease severity ^62^. The latter is consistent with consolidated and efficient use of TMPRSS2 in environments where ACE2-Solute carrier complexes are enriched primarily in the upper respiratory tract (ACE2-SLC6A20) and gastrointestinal tract (ACE2-SLC6A19) and to a lesser extent the lower respiratory tract.

To conclude, this study demonstrates that S FCS cleavage regulation drives the SARS-CoV-2 shift from a pandemic to endemic pathogen. Curiously it is the optimisation of the FCS to remain competitive in one tissue that then culminates in opportunity costs through multifactorial attenuation elsewhere. Continued evolution within this pathway—evidenced by both JN.1’s use of TMPRSS2 and the continued appearance of potential FCS-modulating substitutions including K679R in circulating XFG and BA.3.2.2 lineages—suggests ongoing selection at the FCS locus even during endemic circulation. These findings are important for understanding SARS-CoV-2 from pandemic to endemic stages over the last 6 years and further highlight this consolidated trajectory. Finally, this work provides resources and approaches that can rapidly resolve Omicron lineage phenotypes using clinical material (nasopharyngeal swabs) and thus provides a means for rapid phenotypic tracking of viral evolution and preparation for potential resurgence of variants that may revert to targeting the lower respiratory tract and sustaining greater levels of disease severity.

## Supporting information

Raw data for all figures

## Acknowledgements

This study was supported by NSW Health (Covid-19 response grant F.B., S.G.T, W.R. & J.K.; Prevention Research Support Program, Stream 2: Infectious disease capability, preparedness and response. J.K., S.G.T., R.R., V.S.). FastGrants, an Investigators in the Pathogenesis of Infectious Disease Awards from the Burroughs Wellcome Fund (D.V.). D.V. is an investigator of the Howard Hughes Medical Institute and the Hans Neurath Endowed Chair in Biochemistry at the University of Washington.

## Methods

### Cell culture

HEK293T cells (Thermo Fisher, R70007), HAT-24^2^ (derived from HEK293T cells), IGROV-1 (Sigma, scc203), VeroE6 cells (ATCC CRL-1586) and their derivatives (VeroE6-TMPRSS2^63^, VeroE6-ACE2-TMPRSS2, VeroE6-C4-ACE2-TMPRSS2, VeroE6-TMPRSS2-ACE2+SLC6A19 (TASL-19), VeroE6-TMPRSS2-ACE2+SLC6A20 (TASL-20) and Tet-TASL-19) and A549 cells (Sigma; 86012804-1VL) and their derivatives (AT-T, AT-AT, AT-AT-19, AT-AT-19^3M^ and AT-CT-19) were maintained in Dulbecco’s Modified Eagle Medium (DMEM; Gibco, 11995073) with 10% fetal bovine serum (FBS) (Sigma; F7524). VeroE6 and A549 derived cell lines transduced with TetOne-TMPRSS2 lentivirus were cultured in DMEM supplemented with 10% FBS and either 10 µg/mL or 1 µg/mL puromycin (Sigma; P8833) respectively, until clonally isolated by flow cytometry. For experimental use, VeroE6 and A549 cell lines and their derivatives were plated and cultured in Minimum Essential Medium (MEM; Gibco, 10370-021) supplemented with 2% FBS and 1% L-glutamine (Gibco; 25030-081), while HEK293T and HAT-24 cells used DMEM with 5% FBS. Calu3 cells were maintained in DMEM/F12 media (Gibco;12634010) supplemented with 10% FBS, 1% non-essential amino acids (Sigma, M7145) and 1% Glutamax (Gibco; 35050061). Human nasal airway epithelial cells (NECs) were obtained from brushing the nasal inferior turbinate of eight children. All participants and/or their carers provided written informed consent, with study approval by the Sydney Children’s Hospital Ethics Review Board (HREC/16/SCHN/120). Cells were grown on 6.5 mm Transwell inserts (Corning) pre-coated with PureCol-S collagen type I (Advanced BioMatrix). The cells were incubated (37°C and 5% CO_2_) until confluency in PneumaCult™-ExPlus media (Stemcell Technologies) for 4-7 days before being switched to ALI-culture conditions by removing the apical media and feeding the basal side with PneumaCult™ ALI medium (Stemcell Technologies). The cultures were incubated for 3-4 weeks to achieve mucociliary differentiation, evidenced by the presence of mucus and beating cilia. All stable cell lines used herein were only cultured and used within a range of 20 passages. Each cell line was tested and found negative for mycoplasma (Mycoplasma Testing Facility, UNSW).

### Generation of stable cell lines

For generating VeroE6-TMPRSS2 cells that stably express mutant C4-ACE2, expression plasmid pRRLsinPPT.CMV.GFP.WPRE^2^ was first modified to carry a multiple cloning site (MCS) at the 3’ end of GFP open reading frame (ORF) using Age1 and Sal1 cut sites. The mutant form of hACE2 (C4-ACE2) carrying mutations described previously by Heurich et al ^29^ was synthesized as a synthetic gBlock (Integrated DNA Technologies) and shuttled into the above plasmid using Xba1/Xho1 cut sites thus replacing the GFP ORF with C4-ACE2.

For generating VeroE6-ACE2+SLC6A19 (TASL-19) and VeroE6-ACE2+SLC6A20 (TASL-20) cell lines, expression plasmid pRRLsinPPT.CMV.ACE2.WPRE^64^ was first modified to remove the stop codon at 3’end of ACE2 ORF. A synthetic gBlock with P2A and a MCS carrying Nhe1 cut site was then synthesized (Integrated DNA Technologies) and inserted into the above plasmid at the 3’end of ACE2 to generate ppt-ACE2 MCS plasmid. SLC6A19 (Addgene #161397) and SLC6A20 (Addgene #161409) ORFs were amplified with forward and reverse primers containing Nhe1 and Xho1 cut sites respectively. The SLC6A19 mutant was generated using the same ORF as Addgene #161397, with the exception of inclusions of mutations at F141S, S206V and Y209C. The amplicons were inserted into ppt-ACE2 MCS plasmid using Nhe1/Xho1 cut sites to generate bicistronic plasmids ppt-ACE2-P2A-SLC6A19 and ppt-ACE2-P2A-SLC6A20. A further Collectrin control plasmid was developed by inserting a codon optimised Collectrin ORF fused to P2A (Integrated DNA Technologies) and then ligated over the ACE2-P2A ORF using XbaI and NheI.

For generating the base VeroE6 and A549 TMPRSS2 cells (VeroE6-Tet-TMPRSS2 and AT-T, respectively), the expression plasmid pLVX Tet-One Puro (Clontech) was modified to have a MCS at the 3’end of Tet responsive promoter TRE3GS using EcoR1/BamH1 cut sites. A codon-optimised TMPRSS2 ORF was amplified from a synthetic gBlock (Integrated DNA Technologies) and shuttled into pLVX Tet-One Puro-MCS expression plasmid using Not1/Xho1 sites to generate into pLVX Tet-One Puro-TMPRSS2 plasmid. All plasmid sequences were validated by Nanopore Sequencing with the Rapid Barcoding Kit 96 (Oxford Nanopore Technologies) using the manufacturer’s protocol. The sequencing data was exported as FASTQ files and analyzed using Geneious Prime (v22.2). Open reading frames were then verified using Sanger Sequencing.

Cells expressing ACE2, C4-ACE2, inducible TMPRSS2 and bicistronic ACE2-solute carriers were generated by lentiviral transductions as previously described ^2^. Briefly, lentiviral particles expressing the above genes were produced by co-transfecting expression plasmids individually with a 2nd generation lentiviral packaging construct psPAX2 (courtesy of Dr Didier Trono through NIH AIDS repository) and VSV-G plasmid pMD2.G (Addgene #12259) in HEK293T producer cells using polyethyleneimine as previously described^65^. Virus supernatant was collected 72 h post-transfection, pre-cleared of cellular debris and centrifuged at 28,000 × g for 90 min at 4°C to generate concentrated virus stocks. Lentiviral transductions were then performed on VeroE6-TMPRSS2 cells to generate VeroE6-ACE2-TMPRSS2, TASL-19, TASL-20, on VeroE6 cells to generate Tet-TASL-19, and A549 cells to generate AT-T, AT-AT, AT-AT-19, AT-AT-19^3M^ and AT-CT-19. The expression of TMPRSS2 across the inducible cell lines was stimulated by application of 100 ng/mL of Doxycycline (Sigma, D9891) for 24 h prior to viral exposure, as per the manufacturer’s instructions (CloneTech). Initial screening of A549 clones and selection of clone 8 for further gene transductions was based on expression of high levels of TMPRSS2 following overnight 100 ng/mL Doxycycline stimulation. No such screening selection was made for the VT-T cell line, with a second gene transduction following immediately after 7 days of puromycin selection to create the final Tet-TASL-19 cell line.

### Flow cytometry profiling of cells for ACE2 and TMPRSS2 receptors

To visualise the presence and/or concentration of SARS-CoV-2 entry factors within the desired cell lines, 2 x 10^5^ cells (with or without 24 h doxycycline incubation) were washed with chilled FACS buffer (DPBS containing 1mM EDTA and 1% Human serum) and stained for ACE2 (Human ACE2 APC-conjugated antibody; R&D Systems, FAB933A) and TMPRSS2 (Human TMPRSS2 PE-conjugated antibody; Biolegends, 378402) for 30min on ice in the dark. Cells were washed twice with FACS buffer followed by fixation with 4% paraformaldehyde (final) for 10min, before resuspension in FACS buffer and acquisition by a BD LSRFortessa^TM^ flow cytometer (BD Biosciences). Parental cell lines and/or unstimulated concentrations of the inducible cell line were used as controls.

Gating strategy: the whole cell population was first defined by FSC/SSC analysis, followed by single cell identification by FSC-H vs FSC-A analysis, and finally respective cell populations were defined as ACE2 and/or TMPRSS2 +/- by quadrant gate analysis. Quadrant gates were set based on standard cell controls per replicate experiment. A minimum of 10,000 events were acquired within this established flow cytometry gate for each sample, followed by analysis of the flow cytometry standard (FCS) 3.0 files using FlowJo analysis software (v10.8.0, BD Biosciences).

### Generation of pseudovirus particles and cell entry studies

The generation of vesicular stomatitis virus-based pseudovirus particles bearing SARS-CoV-2 S proteins and cell entry studies were conducted in accordance to previously published protocol^66^. At 24 h post transfection, HEK293T cells expressing the respective S protein, VSV-G or dsRed (negative control) were inoculated with VSV-G-trans complemented VSV∗ΔG(FLuc) (kindly provided by Gert Zimmer) ^67^ and incubated for 1 h at 37°C and 5% CO_2_. Subsequently, the inoculum was aspirated and the cells were washed with PBS before cell culture medium with anti-VSV-G antibody (culture supernatant from I1-hybridoma cells; ATCC no. CRL-2700) was added. Of note, cells expressing VSV-G received cell culture medium without antibody. At 16–18 h post inoculation, the culture supernatants were collected, centrifuged to remove cellular debris (4,000 × g, 10 min), and clarified supernatants were aliquoted and stored at −80°C until further use. For cell entry studies, target cells were grown in 96-well plates to 50–90% confluence before they were inoculated with identical volumes of the respective pseudovirus particles and incubated for 16–18 h at 37°C and 5% CO_2_. Then, cell entry efficiency was assessed through measurement of virus-encoded luciferase activity in cell lysates. Cells were lysed with PBS containing 0.5% Tergitol (Carl Roth) for 30min at RT, before transferring to white 96-well plates. Samples were mixed with luciferase substrate (Beetle-Juice, PJK) and luminescence was subsequently measured with a Hidex Sense plate luminometer (Hidex).

### Expression and analysis of amino acid transport in SLC6A19 and related mutants in Xenopus laevis oocytes

Holding of *X.laevis* frogs (purchased from Nasco) and the surgical procedure to remove parts of the ovary were approved by the Animal experimentation ethics committee of the Australian National University (Protocol #A2023/24). All procedures were carried out in accordance with the recommendations of the Australian code for the care and use of animals for scientific purposes. *X. laevis* oocytes were isolated and maintained as described previously^68^. Selected oocytes were injected with cRNA (12.5 ng each) encoding ACE2 and SLC6A19 and incubated for up to 5 days^40^. The cRNAs were generated using an mMESSAGE mMACHINE T7 Transcription Kit (Thermo Fisher). Mutants of SLC6A19 were generated by site-directed mutagenesis using QuikChange II site directed mutagenesis kit (Agilent). All constructs were confirmed by Sanger sequencing (Biomolecular Resource Facility, Australian National University). Subsequently, uptake experiments were performed using ND96 buffer (96mM NaCl, 2mM KCl, 1.8mM CaCl_2_, 1mM MgCl_2_, 5mM HEPES; titrated with NaOH to pH7.4). For uptake experiments, ND96 was supplemented with 100 μM [^14^C]leucine and incubated with the oocytes for the indicated time. To terminate uptake, oocytes were washed three times with 4 mL of ice cold ND96, transferred to scintillation vials, lysed with 200 μL 10% SDS, and counted for determination of accumulated radioactive.

### Viral isolation, propagation and titration from primary specimens

SARS-CoV-2 variants were isolated from diagnostic respiratory specimens as previously described and through the primary diagnostic provider Douglas Hanly Moir under UNSW ethics IREC5100. Briefly, specimens testing positive for SARS-CoV-2 (RT-qPCR, Seegene Allplex SARS-CoV-2) were sterile-filtered through 0.22 µm column-filters at 10,000 x *g* (10min at 4°C) and serially diluted (1:3) on VeroE6-ACE2-TMPRSS2, C4-VeroE6-TMPRSS2 and/or TASL-19 cells (5 x 10**^3^**cells/well in 384-well plates), followed by incubation for 72 h. Upon confirmation of cytopathic effect by light microscopy, a minimum of 80ul of culture supernatant were added initially to a pellet of TASL-19 cells (1.0 × 10**^6^**cells) for 30 min (37°C with 5% CO_2_) and then subsequently transferred to a 6-well plate with 2 mL of MEM-2%FBS (final). Cells were incubated for 24 to 48 h, or until cytopathic effects had led to loss of >50% of the cell monolayer. The supernatant was collected and cleared by centrifugation (2000 x *g* for 10 min), frozen at -80°C (passage 2), then thawed and titrated on TASL-19 cells to determine median 50% Tissue Culture Infectious Dose (TCID_50_/mL) according to the Spearman-Karber method ^69^. Viral stocks used in this study correspond to passage 3 virus, which were generated by infecting TASL-19 cells at MOI=0.025 and incubating for 24 h before collecting, clearing, and freezing the supernatant as above in 100 µl aliquots. Sequence identity and integrity were confirmed for both passage 1 and passage 3 virus via whole-genome viral sequencing using an amplicon-based Illumina sequencing approach, as previously described ^2^. The latter was also used in parallel for sequencing of primary nasopharyngeal swabs.

Virus titrations were carried out by serially diluting virus stocks (1:5) in MEM-2%FBS before adding to cells in suspension at 5 × 10^3^ cells/well in 384-well plates (with 24 h prior incubation with doxycycline), and then incubating for 48 or 72 h. The cells were then stained live with 5% v/v nuclear dye (Invitrogen, R37605) and whole-well nuclei counts were determined using the IN-Cell Analyzer 2500HS high-content microscope and IN Carta analysis software (Cytiva, USA). Data was normalized to generate sigmoidal dose–response curves (average counts for mock-infected controls = 100%, and average counts for highest viral concentration = 0%) and median 50% nuclei loss NC_50_ values were obtained with GraphPad Prism software ^70^.

### Fluorogenic peptide analysis for TMPRSS2 activity

Cell lines VeroE6, VeroE6-TMPRSS2, VeroE6-ACE2-TMPRSS2 and TASL19 were plated at 2×10^4^ cells/well a 96-well plate for 24 h incubation. Media was subsequently removed and replaced with phenol-red free, FBS-free DMEM media with 100 uM of Boc-QAR-AMC substrate (R&D Systems). Cells were incubated for either 1 h or 3 h at room temperature before fluorescence was measured (Excitation:380nm, Emission:460nm) using the CLARIOstar^PLUS^ microplate reader (BMG Labtech, USA). Each condition was plated as three technical replicates, and the experiment performed in triplicate.

### Viral Plaque assays

Cell lines VeroE6-ACE2-TMPRSS2 and TASL-19 were seeded into 24-well plates at 5 x 10^5^/well in DMEM +10%FBS and incubated for 24 h to achieve a 90–100 % confluency. Cells were inoculated with the desired SARS-CoV-2 variants at 1:10 serial dilutions in DMEM media without FBS, at incubated for 60min (37°C with 5% CO_2_) with gentle agitation every 15 min to ensure even distribution of the virus over the cells. A 2% agarose solution was created in MilliQ water before mixing with an equal volume of DMEM, and subsequently applied to all wells as an overlay media. Cells were incubated for 72 h before a 25min incubation with 4% paraformaldehyde (final). All media and agar were carefully removed, and cell monolayers stained with 1% crystal violet for 10min at room temperature. The solution was then carefully removed and contents allowed to air-dry before size and numbers of plaques were subsequently measured and enumerated.

### Cell-cell transfer assays

Spike constructs for A.2.2 and omicron variants were cloned into a pLVX-zGreen vector, using restriction sites EcoRI and XbaI to create pLVX-spike-A.2.2, pLVX-spike-delta, pLVX-spike-XBB.1.5 and pLVX-spike-KP.3. Spike plasmid constructs were transiently transfected into TASL-19 cells (5 x 10^5^ cells; 500ng plasmid DNA) using Lipofectamine 3000 (Thermo Fisher) following the manufacturer’s protocol. The plasmid pLVX-IRES-ZsGreen was used as a control. After 24 h, cells were stained with 5% v/v nuclear dye (Invitrogen, R37605) and imaged using the IN Cell Analyzer (GE Healthcare, USA) to visualise cell nuclei and fusogenic activity. Images were analysed using the opensource software CellProfiler^TM^ (Broad Institute, USA) to enumerate the nuclei within the spike driven syncytia.

### Assessment of SARS-CoV-2 spike FCS mutations on viral infectivity

A KP.3 R685P mutant was also generated to inactivate the FCS, as described in a previous study ^71^. For Ancestral Clade A.22, the FCS was also inactivated through VeroE6 passaging but sustained a FCS deletion ^679^NSPRRAR^685^ VeroE6 cells (1 x 10^6^) were incubated with a WT KP.3 or Clade A.22 variant (previously isolated and expanded from primary nasal swabs) in MEM + 2%FBS at an MOI of 0.01 following SARS-COV-2 isolate expansion as described previously. After 72 h, the viral supernatant was harvested and centrifuged (1200 x *g*, 10min at 4°C) to remove cell debris, before 0.5mL was heat inactivated and then RNA extracted for whole-genome sequencing as previously described ^2^. A portion of each passage was reseeded into additional VeroE6 cells at a 1:1000 dilution. This process was repeated for a total of six passages. Upon confirmation of a R685P mutation for the KP.3 variant and FCS deletion in Clade A.22, viral titrations were performed alongside a wildtype KP.3 in VeroE6, VeroE6-ACE2-TMPRSS2 with or without 10 µM Nafamostat.

### Detection of catalytic ACE2 in cell supernatants

Soluble ACE2 activity in culture supernatants was measured as previously described^72^.

### Single resolution of known SARS-CoV-2 entry factors with ACE2 associated solute protein carriers

Single cell RNAseq data from nasal ^73^ and ileum ^74^ were obtained from covid19cellatlas.org, while lung data ^75^ was obtained from GEO (GSE171524). These data were analyzed with Seurat V4.3.0 as described previously^76^. Their accompanying metadata, which includes information such as sample ID, sample status, and cluster annotations (cell types), were added to Seurat objects using the “AddMetaData” function. Read counts were normalized using SCTransform, before reanalysis with the standard Seurat workflow of “RunPCA,” “FindNeighbours,” “FindClusters,” and “RunUMAP.” Cluster identities were assigned using published cluster annotations and plots were generated with “DimPlot”, “Featureplot” and “DotPlot”, to illustrate expression of *ACE2*, *TMPRSS2*, *SLC6A19* and *SLC6A20*.

### Soluble Protein expression and purification

The ectodomain of human ACE2 (residues 19-740) was cloned into the pCMV vector with a mu-phosphatase secretion signal peptide and C-terminal His and AviTag by Genscript. The ACE2-C4 mutant was recreated by adding six mutations to the ACE2 CLD: R708A, K713A, R716A, R719A, R721A, and R727A. Two additional mutants were created: ACE2-C4-A containing only R708A, K713A, R716A, R719A, and ACE2-C4-B containing R721A and R727A. All four constructs were expressed in expi293F cells (Thermo Fisher) using transient transfection with Expifectamine (Thermo Fisher) and purified by IMAC using Ni Sepharose Excel resin (Cytiva). Catalytic-dead, disulfide-stabilized TMPRSS2 S441A, T447C ectodomain with an internal enterokinase cleavage site and C-terminal His and AviTag tags was expressed and purified similarly, and activated *in vitro* as described previously^77^. Mature (cleaved) TMPRSS2 was biotinylated with an Avidity BirA Ligase kit (Avidity), then re-purified by gel filtration over a Superdex 200 10/300 column (Cytiva).

### His-Tag Protein Pulldown

Gene-constructs of WT-ACE2, C4-ACE2 mutant, TMPRSS2 and TMPRSS2^S441A^ were subcloned cloned into the pCDNA6 expression vector using cut site enzymes XbaI and XhoI. ACE2 constructs were modified to express a C9-tag on the C-terminal. The TMPRSS2 gene construct was modified to express a 6xhis-tag on its C-terminal end. As above all pCDNA6 plasmid constructs of all genes were verified using Nanopore Sequencing with the Rapid Barcoding Kit 96 (Oxford Nanopore Technologies) using the manufacturer’s protocol. All pCDNA6 were each transiently transfected using PEI into HEK293-T cells with pCDNA6-TMPRSS2-histag as previously described^59^. Transient transfections used a total of 5μg DNA with 5 million cells to ensure there was enough material for later pulldown experiments. The ACE2:TMPRSS2 DNA ratio was 2:1, with empty pCDNA6 being used as a DNA filler. Transfected cells incubated at 37 C for 48 hrs. After incubation, cells were harvested and washed with PBS.

Transfected cells were lysed in RIPA buffer (Thermo Fisher) with EDTA-free Protease Inhibitors (Roche). Lysate was applied on Ni-NTA Magnetic Beads (S1423S, New England Biolabs) using manufacturers protocol. A fraction of raw lysate was saved for blotting. After samples were eluted from the magnetic beads, they were diluted in NuPAGE LDS Sample Buffer (4x) and NuPAGE Sample Reducing Agent (10X) (Thermo Fisher), then samples were boiled at 100°C for 5 mins. Samples underwent SDS-PAGE and Western Blotting with conditions described previously. Primary antibodies used were anti-C9 antibody (1:500, Santa Cruz, sc-57432) and anti-SLC6A19 antibody (1:2000, HPA043207-100UL, Sigma-Aldrich). Anti-C9 blots were incubated in goat anti-mouse HRP conjugate (1:10,000, Bio Rad, 1721011). Anti-SLC6A19 blots were incubated in donkey anti-rabbit HRP conjugate (1:10,000, A16023, ThermoFisher). Visualization was performed using a G:Box (Syngene).

### Immunoprecipitation of soluble proteins

To prepare TMPRSS2 affinity resin, 1.4 mg of superparamagnetic Streptavidin beads (MagReSyn) was equilibrated in 50 mM HEPES pH 7.4 and 150 mM NaCl (HBS 50/150), then incubated with 130 µg biotinylated TMPRSS2 for 1 hour at 4°C with gentle agitation. The flow-through contained approximately 14 µg by A280 indicating saturation by TMPRSS2. Using a magnetic stand, the resin was washed with 500 µl (>>100 CVs) of HBS with 500 mM NaCl (HBS 50/500) and then four additional times with HBS 50/150.

The immobilized TMPRSS2 resin was used for IPs with all four WT or mutant ACE2 proteins. Bonafide IPs employed 300 µg resin and 160 µg ACE2 in a total volume of 200 µL HBS 50/150. Mock IPs were also conducted using 300 µg of bare Streptavidin resin incubated with 320 µg wild-type or mutant ACE2 in 150 µL HBS 50/150. After incubating overnight at 4°C with agitation, the flow-through solutions were removed, and the beads were washed once with 150 µL HBS 50/500 and then four additional times with 150 µL HBS 50/150. Low-pH elution was conducted by incubating resin in 90 µL 100 mM glycine pH 2.2 for 15 minutes, followed by a rinse with 60 µL additional glycine, and eluent neutralization with 8 µL of 1N NaOH. SDS-PAGE samples for each fraction and ACE2 and TMPRSS2 standards (100 ng each) were prepared in 1x NuPAGE LDS (Invitrogen) and run on a 7.5% PAGE gel. The gel was stained with Sypro Ruby (Invitrogen) using the low-background “rapid protocol” and imaged on a GelDoc (BioRad).

### Mass Photometry

ACE2 mutants were prepared as 100 nM stocks in HBS 50/150 and analyzed on a TwoMP mass photometer (Refyne Ltd), using the drop dilution method to obtain tractable event rates. Glass microscope slides were washed extensively with isopropanol and ultrapure water and dried on a nitrogen gas line, and tissue culture gaskets were used to create wells on the slide. Within a well, a 2 µL ACE2 was added to a 18 µL droplet of HBS, and the TwoMP was focused and used to collect movie data. The true concentration of ACE2 was thus near 10 nM. DiscoverMP analysis software (Refyne Ltd) was used to extract and quantify photometry events, apply calibration using in-house protein standards, and plot count data.

### Biolayer-interferometry

Biotinylated TMPRSS2 or Ancestral SARS-CoV-2 Wu1 RBD was loaded to 1 nm shift on Streptavidin tips hydrated in kinetics buffer and then dipped into 2 µM ACE2 or mutant. All steps were performed at 30°C and 1,000 RPM, and association and dissociation phases were monitored for 500s. Baseline subtracted data were plotted with matplotlib^78^.

### SDS-Page and Western Blotting

Cell pellets from uninfected cells and SARS-CoV-2 viral products (as detailed below) were lysed in RIPA buffer (Thermo Fisher) with EDTA-free Protease Inhibitors (Roche). Alternatively, viral supernatants were first centrifuged to remove all cell debris (1000 x *g*, 5min at 4°C) before proceeding with lysis of virions. All lysate and elution samples were subsequently diluted in NuPAGE LDS Sample Buffer (4x) and NuPAGE Sample Reducing Agent (10X) (Thermo Fisher), and incubated at 100°C for 5min. A gel electrophoresis tank (SureLock Tandem Midi gel tank, Thermo Fisher) was prepared with a NuPAGETM Bis-Tris Protein Gel (Thermo Fisher), samples were loaded alongside a protein ladder (Bio-Rad, Cat#1610375) and run at 200V for 35min at room temperature. The gel subsequently underwent a wet transfer onto an immunobilin PVDF transfer membrane (Sigma) at 30V for 90min at 4°C, followed by application of a blocking reagent (1x PBS + 0.1% Tween20 + 2.5% (w/v) skim milk) for 60min at room temperature and rocking. The primary antibody was applied for overnight incubation (4°C, rocking), and membranes washed three times with 1x PBS + 0.1% Tween20 before the secondary antibody was applied (60min at room temperature, rocking). After a final three washes, transfer membranes were soaked in an ECL mixture (ClarityTM Western ECL Substrate, Bio-Rad) for 5min covered in foil before immediately using chemiluminescent imaging (ChemiDocTM Touch Imaging System, Bio-Rad).

Primary antibodies were anti-rhodopsin antibody (1:500, Santa Cruz, sc-57432), anti-SLC6A19 antibody (1:2000, HPA043207-100UL, Sigma-Aldrich), anti-SLC6A20 antibody (1:2000, PA5-68332, Thermo Fisher), and anti-GAPDH antibody (1:4000, ab8245, Abcam). Secondary antibodies included donkey anti-mouse HRP conjugate (1:5000, A16011, Thermo Fisher) and donkey anti-rabbit HRP conjugate (1:10000, A16023, Thermo Fisher). For samples containing SARS-CoV-2 elements, primary antibodies used were SARS/SARS-CoV-2 Spike Protein S2 antibody (1:2000, MA535946, Thermo Fisher), and either anti-SARS-CoV-2 Nucleoprotein (1:2000, Cellabs (custom order)) or anti-GAPDH antibody, followed by the secondary antibody of donkey anti-mouse HRP conjugate.

### Expansion and Purification of SARS-CoV-2 Viral Products

Cell suspensions of 1 x 10^6^ cells (primary nasal epithelial cell organoids, Calu-3, IGROV-1, VeroE6, VeroE6-TMPRSS2, VeroE6-ACE2-TMPRSS2 and TASL19) were created with the desired SARS-COV-2 variant in MEM+2%FBS at MOI=0.025 (based on the calculated TCID50 of previous stock titration on the TASL-19 line) and incubated for 60min (37°C with 5% CO_2_). Infected cells were pelleted (300 x *g* for 5min) and washed twice with MEM+2%FBS, before a final resuspension and plating into a 6-well plate for a 60min incubation. For Nafamostat treatment, the media was carefully removed and replaced with MEM+2%FBS containing 10 μM Nafamostat final concentration. In all cases, cell incubation continued for 24 h before cells and/or supernatant were subsequently harvested. Supernatant was centrifuged to remove cell debris, while cells were washed in PBS and pelleted. Both sample types were stored at -80 C until required for western blotting.

### K18-ACE2 transgenic mice

The two constructs (WT AC2 and C4-ACE2) were removed from the vector backbone by HpaI plus XbaI double digest and microinjected into C57BL/6 embryos using the approach outlined to produce the original K18-hACE2 mice^79^. Pups derived from microinjected embryos were screened by PCR and DNA sequencing and founder mice backcrossed for two generations with wild-type C57BL/6 mice. All procedures involving WT ACE2 and C4-ACE2 mice were approved by the Sydney Local Health District Animal Welfare Committee (2024-025) and Institutional Biosafety Committee (IBC 20-051). Hemizygous WTACE2 and C4-ACE2 K18-hACE2 mice were bred at the Australian Bioresources facility (Moss Vale, NSW) and transported to the BSL3/PC3 facility at the Centenary Institute for infection. Mice were allowed one week for acclimatisation prior to SARS-CoV-2 infection. Mice were lightly anaesthetised with isoflurane and then intranasally administered with 10^3^ PFU of either Ancestral or KP.3 Omicron. Following infection, mice were weighed and clinically evaluated daily. At day 6 post-infection, mice were humanely euthanised with an overdose of sodium pentobarbitone with lung and brain tissues collected and homogenised for plaque assay quantification as previously described ^80,81^.

### Statistical analysis

Statistical analyses were performed using a combination of GraphPad Prism 9 (version 9.1.2, GraphPad software, USA) or 2026 Originlab Pro (MA, USA). Sigmoidal dose response curves and interpolated NC50 values were determined using Sigmoidal, 4PL model of regression analysis in GraphPad Prism. For statistical significance, the datasets were initially assessed for Gaussian distribution, based on which further analysis was performed. For datasets that followed normal distribution, unpaired t-test was used to compare two groups. Samples without normal distributions were subject to non-parametric Mann-Whitney tests for significance. Details of statistical tests used for different data sets have also been provided in figure legends, with statistical analysis for each figure presented in Supplementary table I.

---

**Figure S1.**
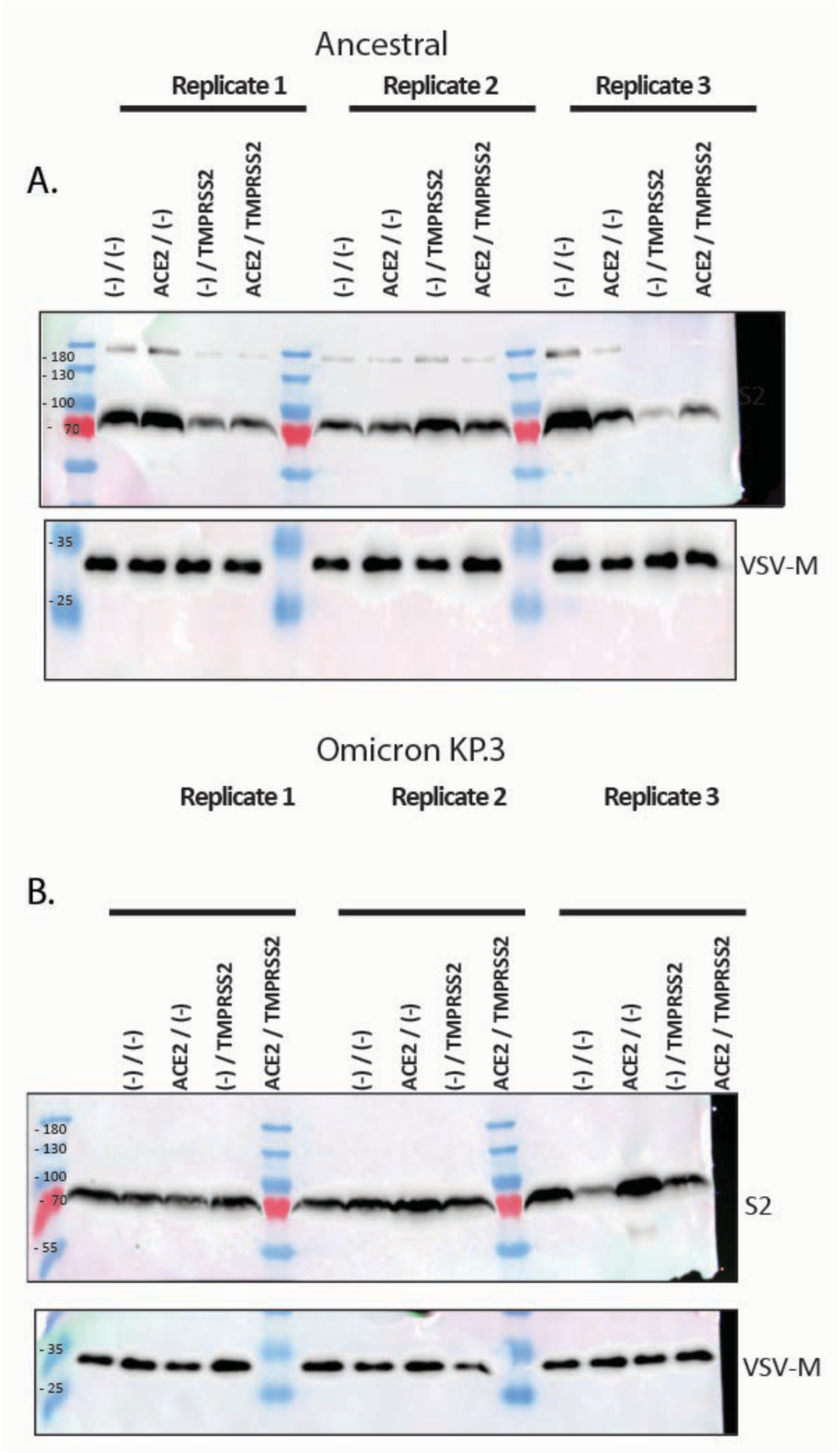
Spike S1/S2 cleavage in representative pre-Omicron and Omicron VSVg pseudotypes. **A. & B.** All receptor constructs in lentiviral constructs used herein were shuttled to the transient mammalian expression vector pcDNA6. Pseudovirus particles were then generated in the presence or absence of various receptor combinations. “(-)/(-)” indicates no receptor control “ACE2/(-)”-WT ACE2, “(-)/TMPRSS2”-TMPRSS2 and “ACE2/TMPRSS2” -combined ACE2 and TMPRSS2 expression during pseudoparticle production in HEK-293T cells. Three days post transfection, pseudoviruses were pelleted via ultracentrifugation for 90’ at 100,000xg through 20 % sucrose cushion. Pellets were then lysed and subjected to Western blotting. Here pseudoviral proteins-VSV-M represents the viral loading control and spike protein is detected for S1/S2 cleavage using the S2 antibody as described herein for primary isolates. **A.** Represents an early circulating Pre-Omicron B-Clade, whilst **B.** represents the more recent Omicron KP.3 lineage. Size of molecular weight markers are presented to the left of Western blots and are in KDa. In Both A. and B. three independent viral pseudotype preparations are presented.

**Figure S2.**
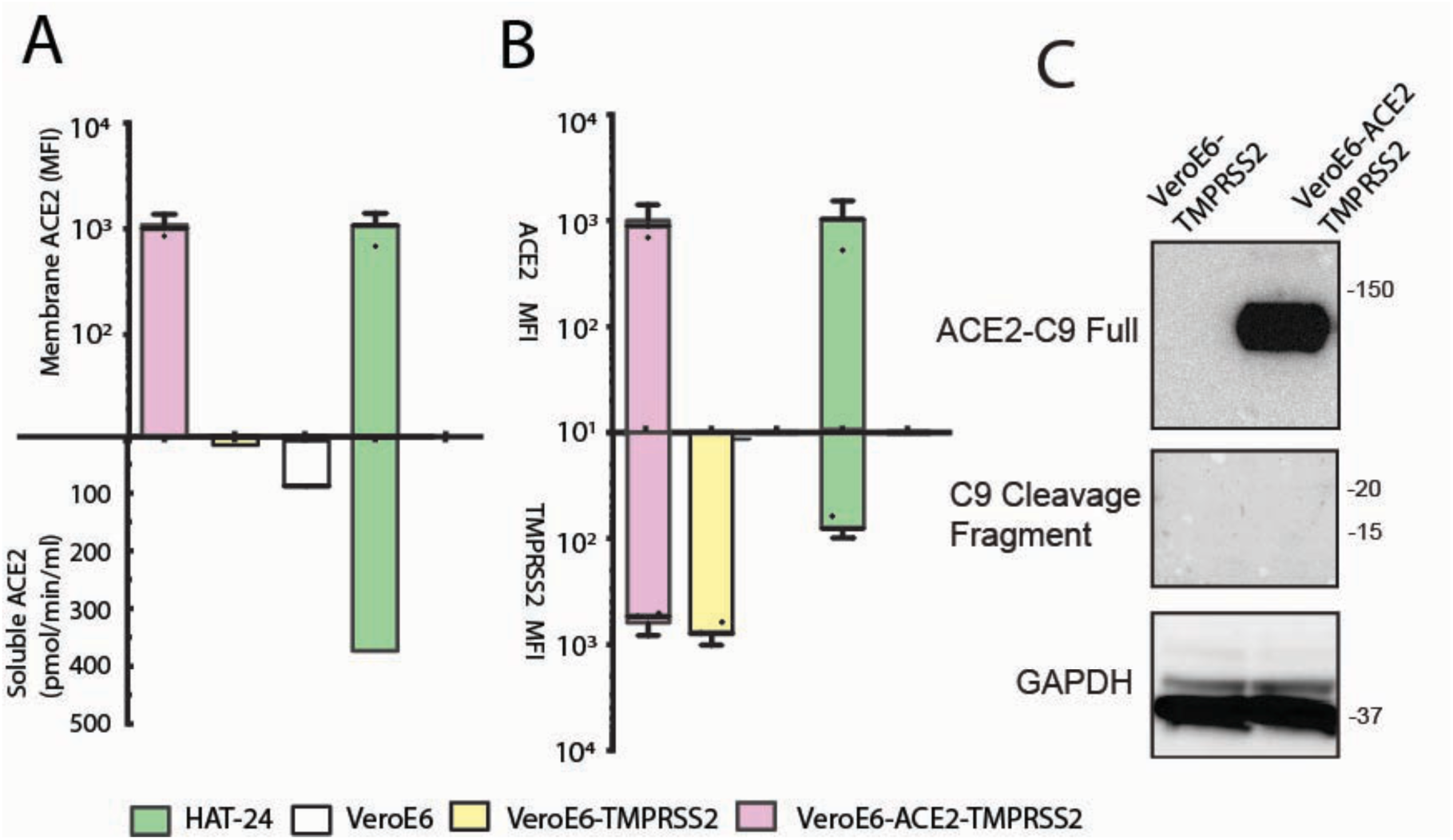
Detection of soluble and membrane bound ACE2 across engineered cell lines. **A.** Membrane and soluble ACE2 levels in engineered cell lines. Membrane ACE2 levels were measured by flow cytometry, mean fluorescence intensity is shown. (Human ACE2 APC-conjugated antibody; R&D Systems, FAB933A) and TMPRSS2 (Human TMPRSS2 PE-conjugated antibody; Biolegends, 378402). The enzymatic activity of soluble ACE2 in the supernatant of 80% confluent cell lines after three days of culture was determined as previously described ^1^. **B.** Ratios of cell surface ACE2 and TMPRSS2 detected via flow cytometry as per A. The HAT-24 line is used as a control, as it is a Hek based cell line that expresses high levels of ACE2^2^ with accompanying ACE2 shedding. **C.** Expression of full length and cleaved ACE2 in VeroE6-TMPRSS2 and VeroE6-ACE2-TMPRSS2 cells determined by immunoblotting with anti-C9 tag antibody relative to GAPDH controls (1:500, Santa Cruz, sc-57432; 1:4000, ab8245, Abcam). Position of molecular weights are to the right and are listed in KDa.

**Figure S3.**
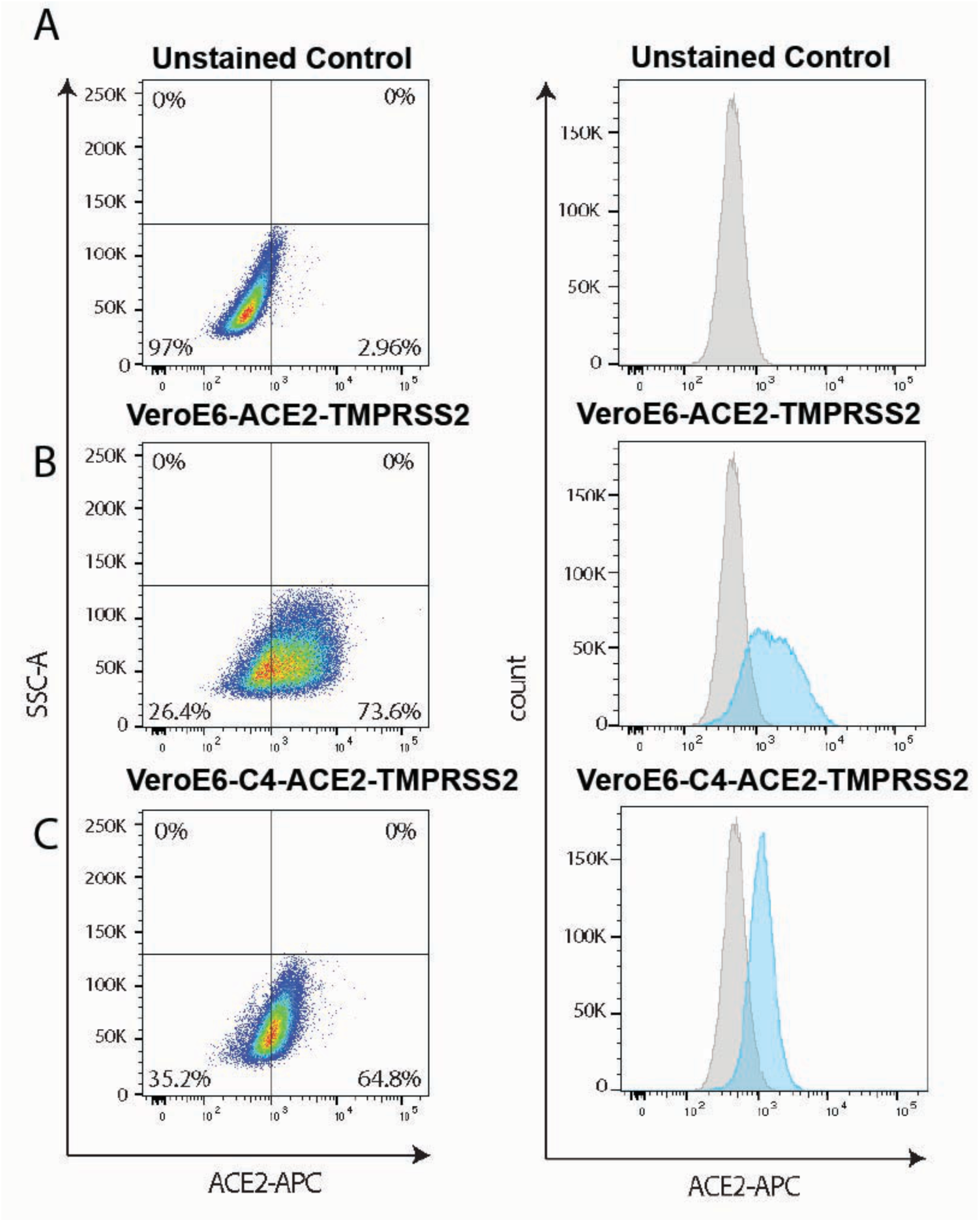
ACE2 surface expression of the VeroE6-TMPRSS2 cell line with WT ACE2 and ACE2 C4 mutant (C4-ACE2). **A.** Unstained control. **B.** VeroE6-ACE2-TMPRSS2 stained for ACE2. **C.** VeroE6-C4-ACE2-TMPRSS2 stained for ACE2. (Human ACE2 APC-conjugated antibody; R&D Systems, FAB933A). Staining outlined in methods. Stains are representative of at least 3 independent flow cytometry stains/acquisitions.

**Figure S4.**
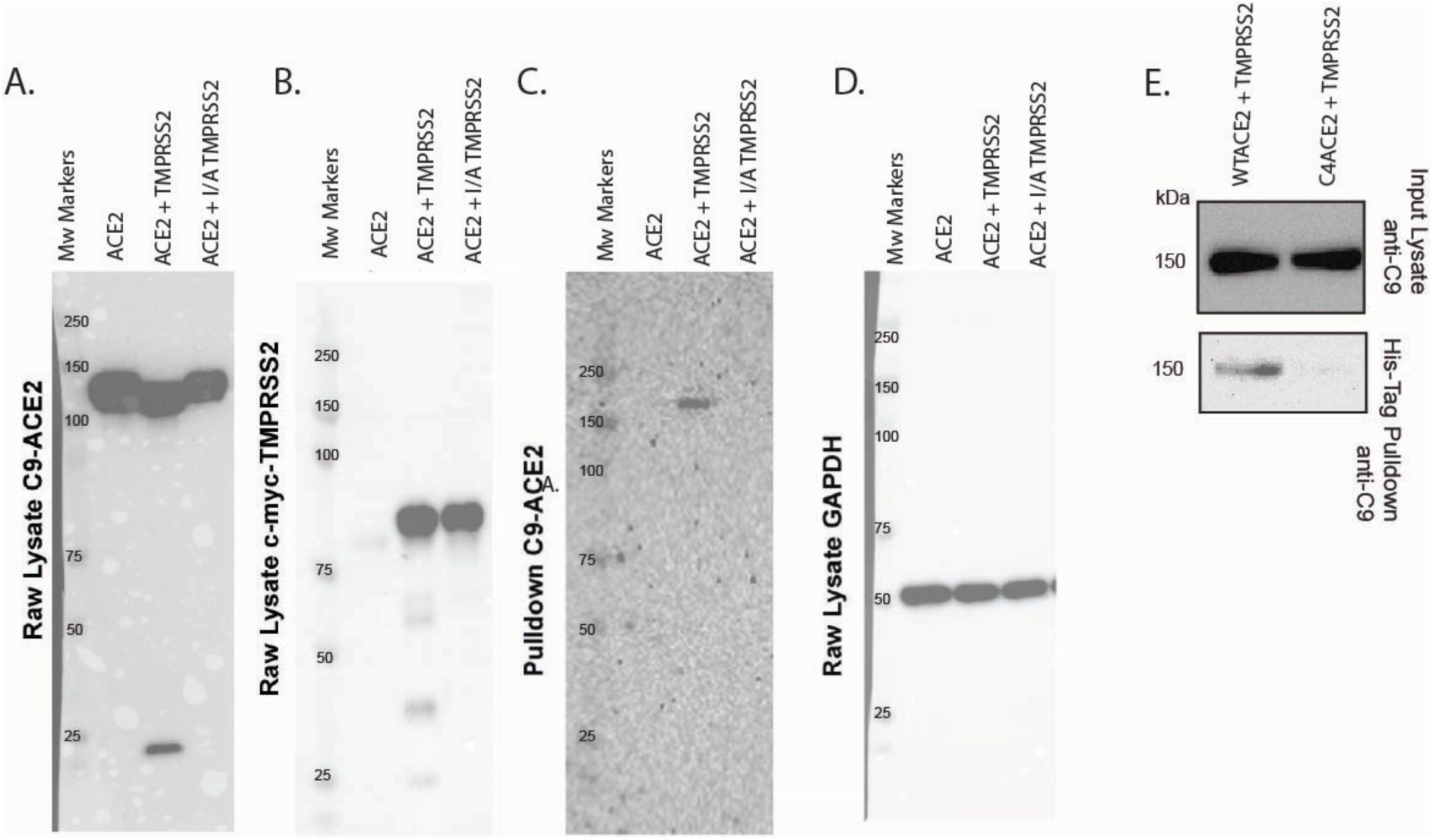
Detection of ACE2-TMPRSS2 complexes in transient expressions. Transient PEI transfection of HEK293T with indicated ACE2 and TMPRSS2 plasmid constructs (all based on the CMV expression vector pCDNA6) at ACE2:TMPRSS2 DNA ratio was 2:1, with empty pCDNA6 being used as a DNA filler DNA when ACE2 is expressed alone. **A.** Detection of ACE2 (anti-C9, Santa Cruz, sc-57432) and **B.** TMPRSS2 (Thermofisher c-Myc Monoclonal Antibody 9E10) in raw lysates in transient transfections of ACE2, ACE2 + TMPRSS2 and ACE2+ catalytic inactive TMPRSS2 (S441A mutant). **C.** TMPRSS2 pull-down of ACE2 WT using Ni-NTA Magnetic Beads (S1423S, New England Biolabs) to capture the His tag at the N-terminus of TMPRSS2 and blotting for ACE2 as in A. **D.** GAPDH control blotting for raw lysates used in A. to C. (Santa Cruz, sc-57432). **E.** TMPRSS2 pull-down of ACE2 WT and ACE2 C4-ACE2 under conditions outlined in C.

**Figure S5.**
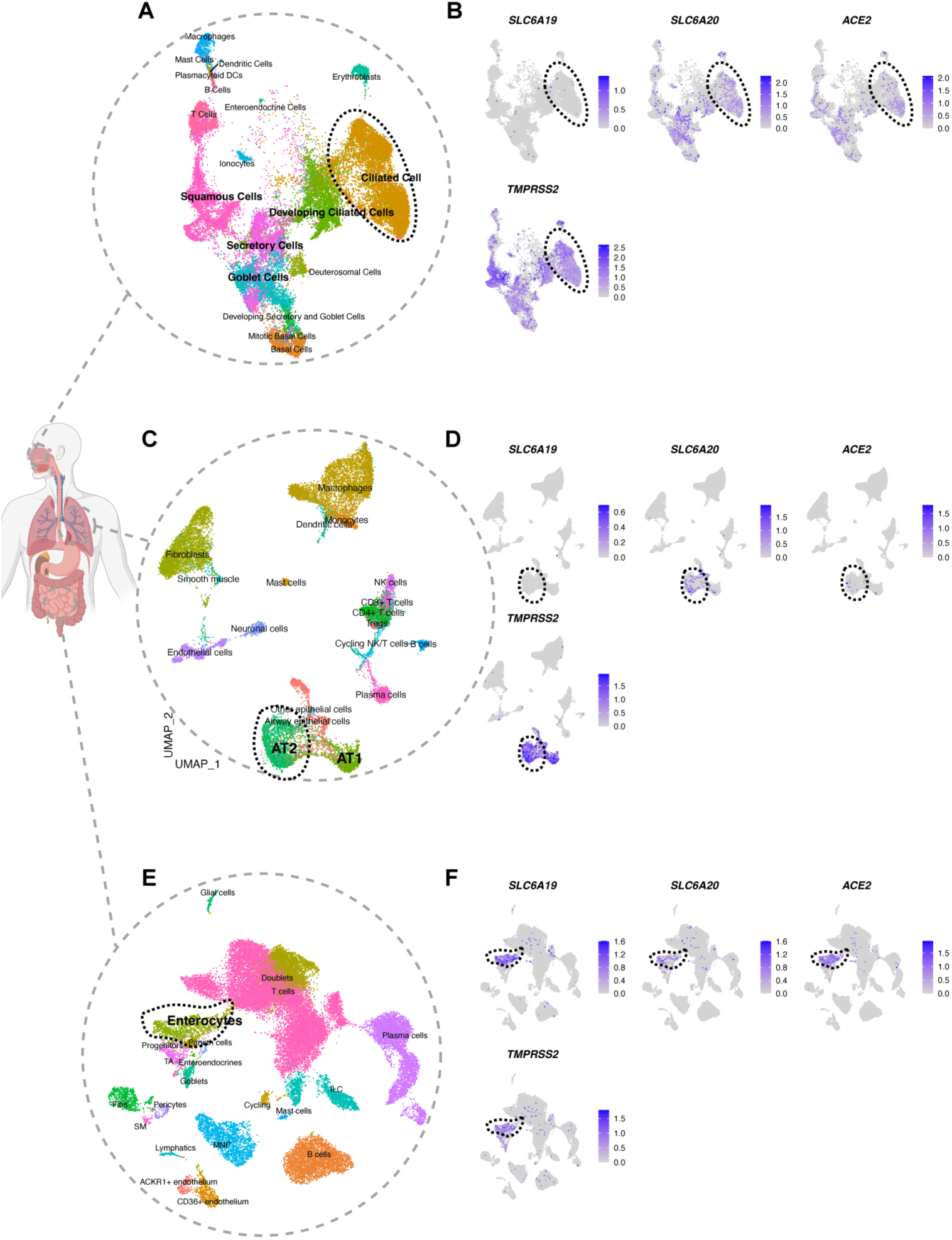
*In vivo* profiling of solute carriers SLC6A19 and SLC6A20 as additional SARS-CoV-2 entry factors alongside known entry factors ACE2 and RSS2. **A.-F.** Uniform manifold approximation and projection (UMAP) of **A.** Nasal Epithelia. **C.** Lung and **E.** Small intestine using single nuclei RNA sequencing. Cell types are color-coded. Target cells within each tissue are highlighted in B. D. & F. in respective tissues, with expression of protein solute carriers SLC6A19 and SLC6A20 alongside the known SARS-CoV-2 receptors ACE2 and TMPRSS2. Of note, ACE2 single cell expression is closely aligned with SLC6A19 and SLC6A20 in the small intestine and nasal cavity respectively. In contrast, the lung is primarily defined by high levels of TMPRSS2 expression in type 2 pneumocytes.

**Figure S6.**
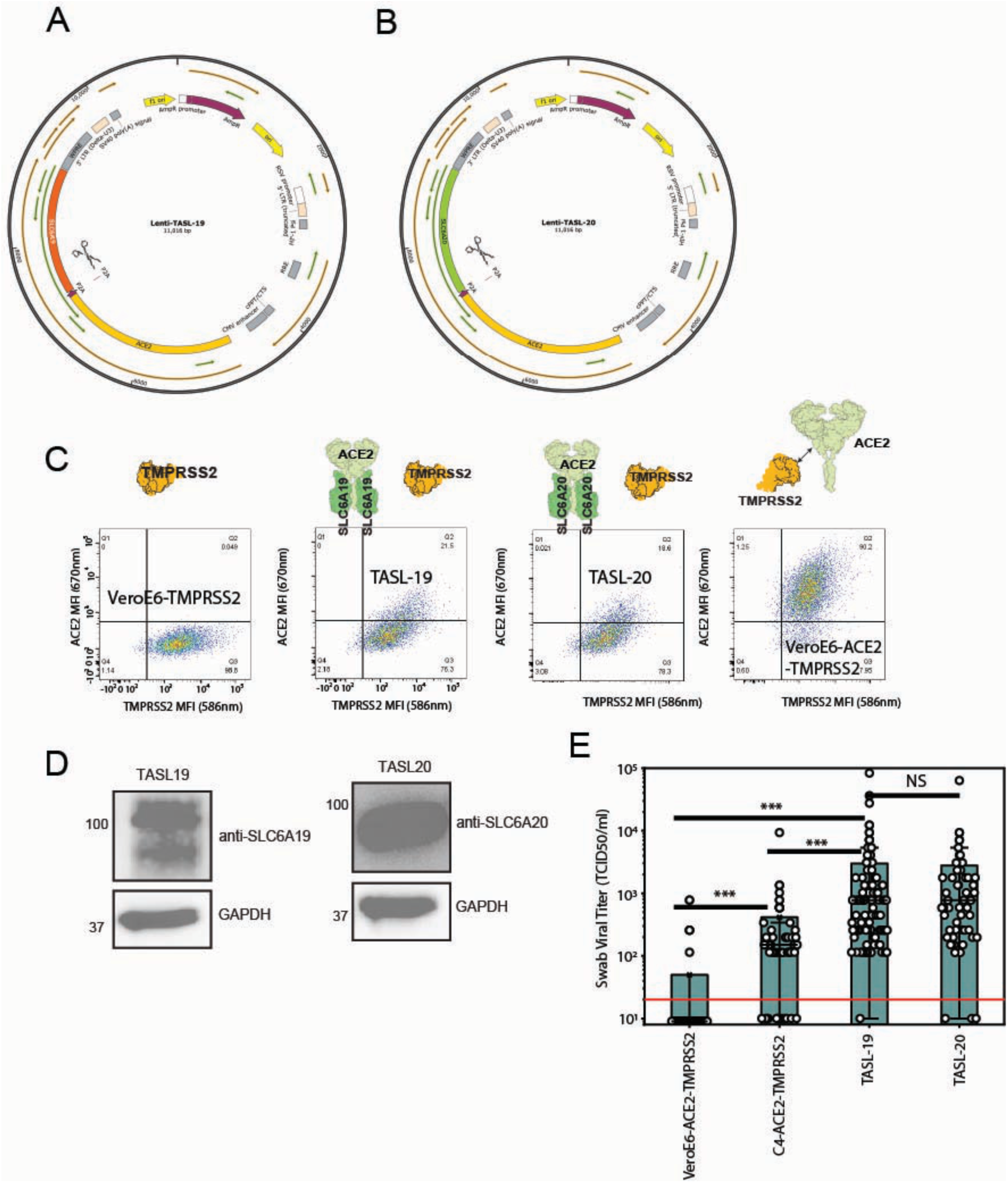
Establishment of the TMPRSS2-ACE2-Solute Carrier Cell lines TASL-19 and TASL-20. **A. & B.** Maps of bicistronic lentiviral constructs to enable equimolar protein expression of ACE2 and A. SLC6A19 or B. SLC6A20. Scissors highlight the P2A cleavage site that liberates ACE2 and either solute carrier. **C.** Development of the TASL-19 and TASL-20 cell lines. Here lentiviral constructs from A & B have been used to genetically modify the VeroE6-T2 cell line with either ACE2 and SLC6A19 or B. SLC6A20. To validate integration and expression, cells were stained using ACE2 and also TMPRSS2 (Human ACE2 APC-conjugated antibody; R&D Systems, FAB933A) and TMPRSS2 (Human TMPRSS2 PE-conjugated antibody; Biolegends, 378402). The parental cell line VeroE6-T2 and the ACE2-VeroE6-T2 are presented as controls. In right histograms data is presented for three independent stains for ACE2 and TMPRSS2. **D.** Given the close proximity of Solute Carriers to the cell membrane, we generated cell lysates from both TASL-19 and TASL-20 clones to confirm expression of each at the protein level. Given their expression is through a bicistronic vector, ACE2 expression alone can act as a surrogate for quantitative solute carrier expression in each cell line**. E.** Initial validation of the TASL-19 and TASL-20 cell line using SARS-CoV-2 positive nasopharyngeal swabs obtained from June to December 2025 when JN.1 sub-lineages were circulating in the Australian community. Each swab serially diluted across each cell line in quadruplicate, scored for cytopathic effects after 72 hours of culture and TCID50 titers then calculated using the Karber method. Red line represents the limit of detection, where negative titers are attributed the titer value of 9 for presentation on the Y-Axis log scale. ***<0.001 Mann-Whitney. NS = Not Significantly different.

**Figure S7.**
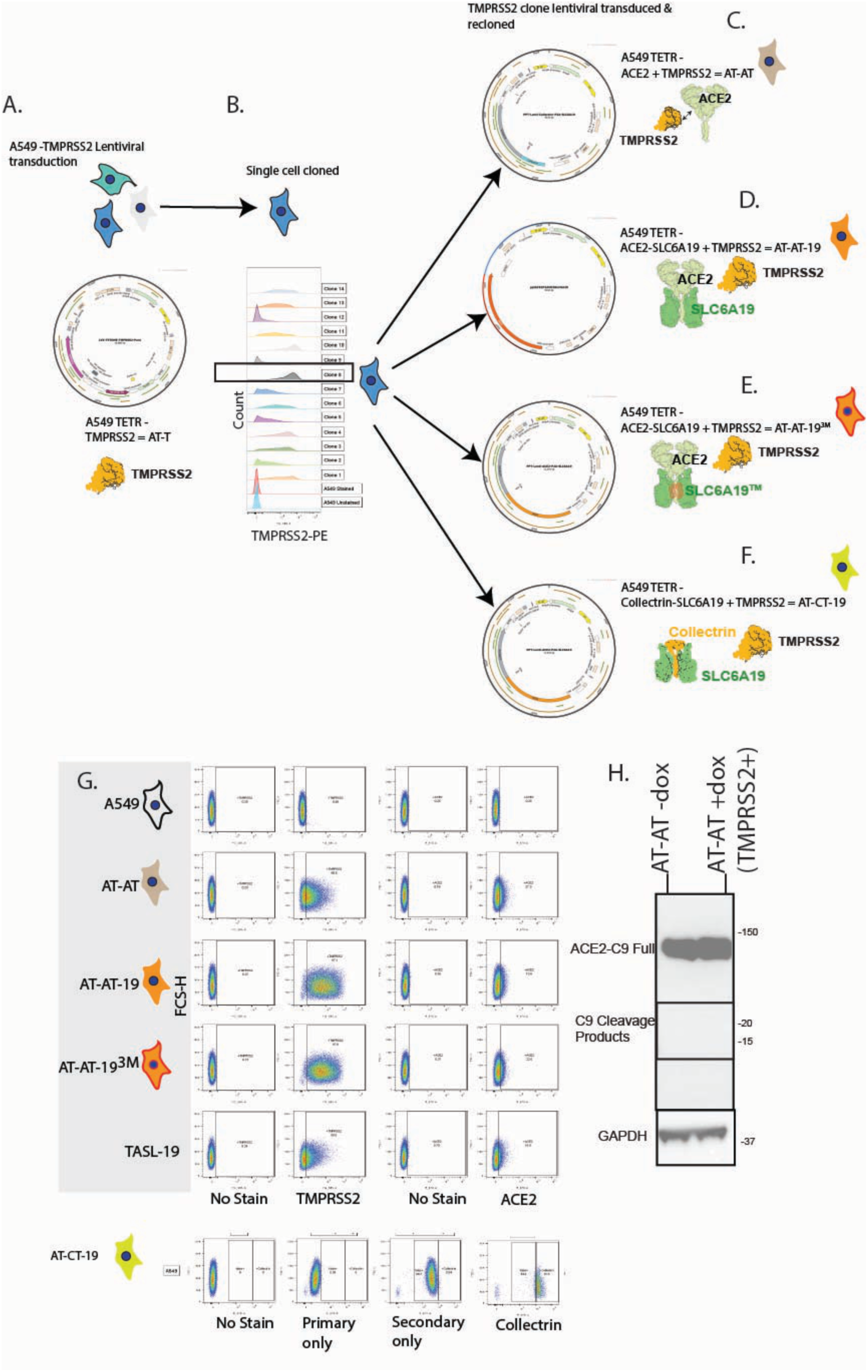
Panel of engineered Lung derived A549 cells. **A.** Strategy to develop A549 cell clones with ACE2, TMPRSS2 and Solute Carriers. TMPRSS2 is introduced under a Tet promoter and maximally induced after 24 hours with 100ng/ml of Doxycycline. **B.** Clones were generated through single cell sorting, stained for TMPRSS2 with clone 8 serving as the template for cells C. to F. **C.-F.** Cells were lentivirally transduced with **C.** ACE2 alone **D.** ACE2 & SLC6A19 using P2A bicistronic lentiviral constructs to enable equimolar protein expression of ACE2 & SLC6A19 **E.** As outlined for D. but with the SLC6A19 triple mutant (SLC6A19^3M^) unable to function as an amino acid transporter. **F.** Collectrin substituting for ACE2 as a chaperone for SLC6A19. This facilitates SLC6A19 surface expression independent of ACE2. **G.** Validation of TMPRSS2, ACE2 and Collectrin expression. Given bicistronic expression vectors, this also acts as a surrogate for SLC6A19. **H.** Lack of ACE2 cleavage with and without TMPRSS2 expression in the AT-AT cell line. Expression of ACE2 in the AT-AT cell line with (+Doxycycline) and without (no Doxycycline) TMPRSS2 expression determined by immunoblotting with anti-C9 tag antibody. Position of molecular weights are to the right and are listed in KDa.

**Figure S8.**
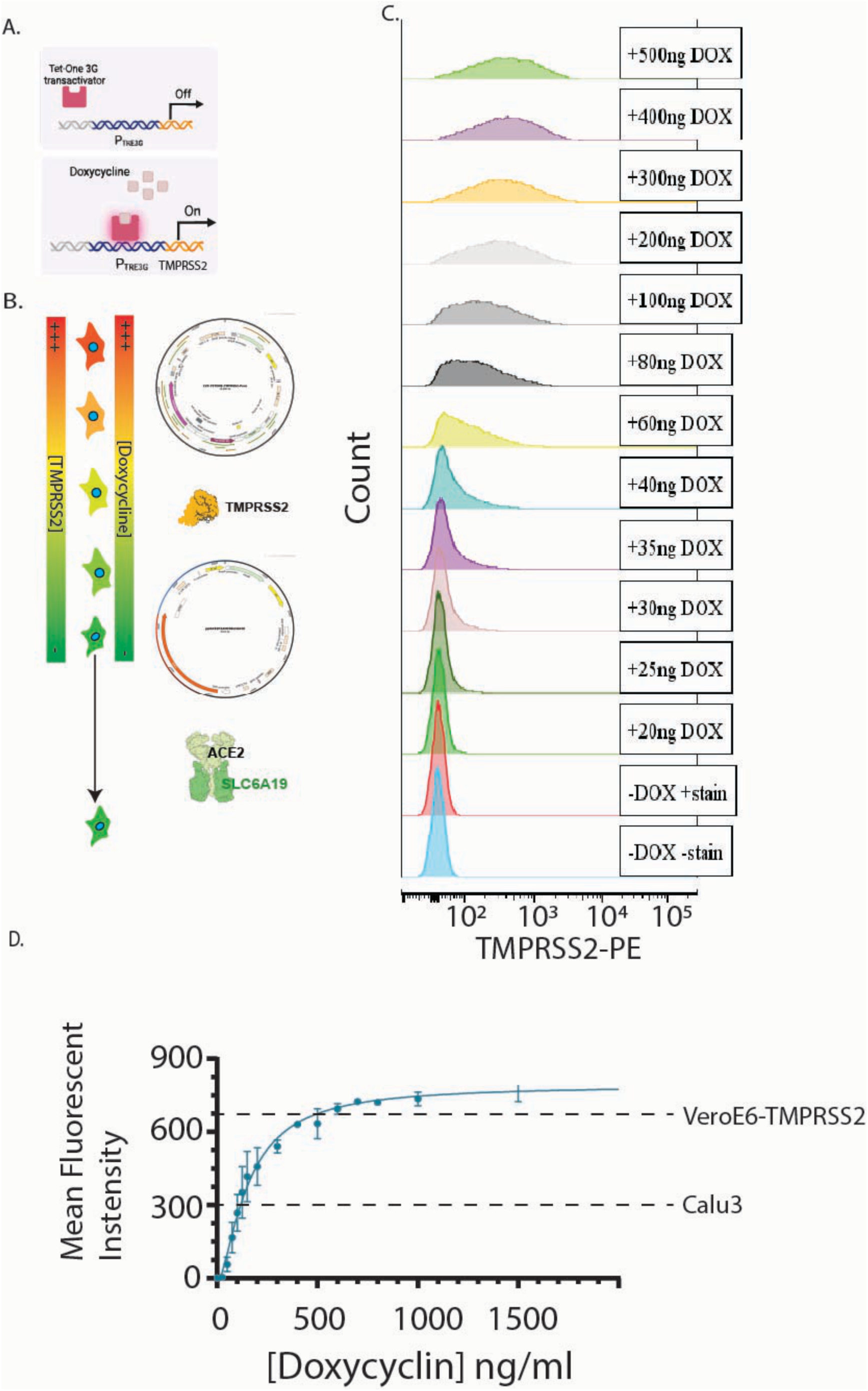
Tet-TMPRSS2 TASL-19 cell line for Doxycyclin regulated expression of TMPRSS2 in the presence of ACE2+SLC6A19. **A.** Schematic of TET promoter based expression. **B.** Generation of the Tet-TASL-19 clonal cell line. Cells were first lentivirally transduced with Tet-TMPRSS2 and clonally sorted. Clones were then selected following 24 hours of 200ng/ml Doxycyclin and flow cytometry as outlined in Fig. S7B. Cells were then lentivirally transduced using the ACE2-P2A-SLC6A19 construct. The latter ensures TMPRSS2 is quarantined from ACE2 based regulation. **C&D.** Final clone expressing ACE2, SLC6A19 and TMPRSS2. Here, tuneable expression of TMPRSS2 is presented as **C.** raw flow cytometry histograms and **D.** aggregate flow cytometry data presented as mean fluorescent intensity with background fluorescence (no staining controls) versus [Doxycyclin]. Values in D. are the mean of three independent Doxycyclin stimulations and stains, with error bars representing standard deviations. Relative expression of TMPRSS2 for VeroE6-TMPRSS2 and unmodified Calu3 are indicated as dash lines with respect to MFI.

**Figure S9.**
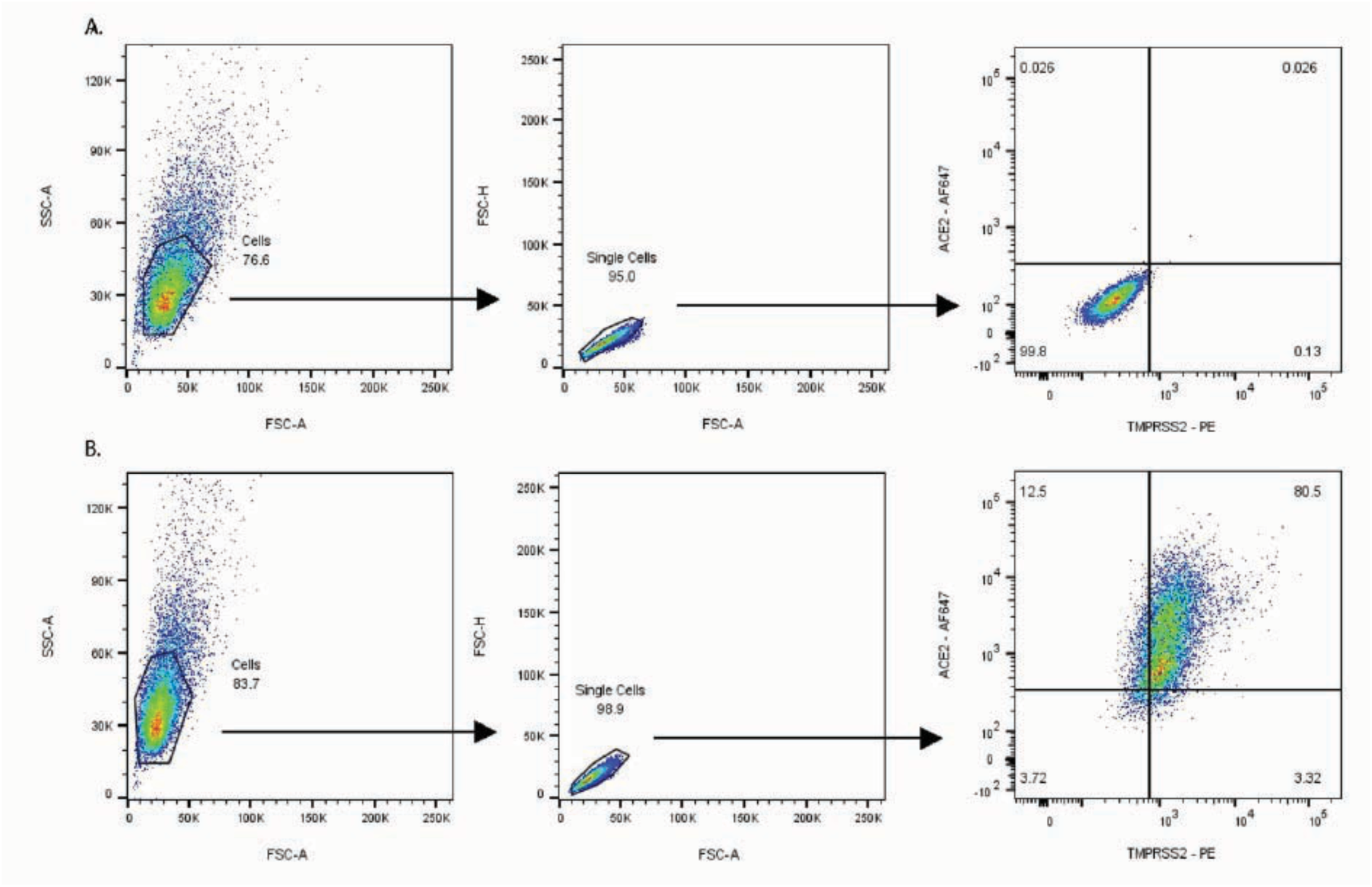
Gating strategy for Flow cytometry plots that are outlined herein. **A & B.** Cell suspensions are generated and stained as outlined in the methods. **A.** Unmodified VeroE6 cells are presented with the left panel establishing gates for forward scatter and side scatter, middle panel for single cells and right panel the resulting co-stain for ACE2 using murine anti-ACE2 conjugated to Alexa-Fluor 647 (Y-Axis) and murine anti-TMPRSS2 conjugated to phycoerythrin. **B.** Represents VeroE6 clones engineered to express human ACE2 and TMPRSS2 (VeroE6-ACE^-^TMPRSS2). Gating strategy and is outlined as in A. In both A&B and minimum of 10,000 events are acquired following establishment of the two gates in the left and central panels.

